# Flexible phosphorus allocation to the phosphoproteome underpins diatom regulatory adaptations to fluctuating phosphate supply

**DOI:** 10.64898/2026.11.11.763867

**Authors:** Yasmin Meeda, Yousef Touhami, Susan Wharam, GE Jiayou, Ellen L. Harrison, Qiong Zhang, Kate J. Heesom, Philip A. Lewis, Adam Monier, Glen L. Wheeler, Katherine E. Helliwell

## Abstract

Diatoms are major drivers of marine phosphorus (P) cycling and are highly adapted to cope with variable P supply. P-starved diatoms can sense increases in P availability within seconds, accumulating this limiting resource through luxury polyphosphate (polyP) storage. Yet, how diatoms coordinate such responses is unknown. Protein phosphorylation is a core mechanism for environmental signal transduction in eukaryotes, but how cells balance investment in protein phosphorylation against competing physiological demands during P limitation is poorly understood. Here, we characterise system-wide remodelling of the *Phaeodactylum tricornutum* phosphoproteome in response to P deficiency and subsequent resupply. We identify widespread proteome dephosphorylation across 44% of phosphosites, revealing a previously unrecognised P-sparing strategy in marine phytoplankton. Following P resupply, diatoms orchestrate extensive phosphoproteome reprogramming within minutes. We characterise previously unrecognized components for early recovery from P limitation, including a novel environmentally prevalent intracellular H⁺/phosphate transporter strongly conserved in diverse diatoms. We also identify a vacuolar Ca²⁺-dependent protein kinase (CDPK) associated with polyP-storage granules. Mutant and inhibitor analyses demonstrate a role for Ca^2+^- and CDPK-signalling in the regulation of luxury polyP accumulation. Our findings reveal that recovery from P starvation is mediated by rapid phosphoregulatory control of intracellular P trafficking, organellar ATP generation coupled with luxury P storage, enabling diatoms to exploit newly available P and thrive in fluctuating P regimes. Collectively, our study uncovers the phosphoproteome as a flexible intracellular P reservoir and establishes mechanistic links between intracellular P budgeting, signalling and the marine P cycle.

## INTRODUCTION

Diatoms are abundant primary producers, responsible for ∼20% of global carbon fixation [1–3]. Thriving in coastal and upwelling systems, they are often first to respond to nutrient influxes, dominating early bloom formation [4]. Phosphorus (P) is a crucial macronutrient frequently found in limiting supply in aquatic ecosystems [5–8]. Yet, river run-off, upwelling and turbulent mixing can cause rapid elevations in P availability [9, 10]. Diatoms sense such shifts within seconds [11], leading to metabolic responses [12] and gene expression changes within hours to days [13]. However, despite the likely importance of nutrient signalling processes in driving bloom formation, the sequence of events from initial sensing of phosphate to coordinated downstream responses is a major black box. Whilst ‘omics studies have been instrumental for deducing how diatoms cope with P starvation and recover following resupply [12–14], regulatory responses at the post-translational level (e.g. protein phosphorylation), which likely play a key role in governing such adaptations, remain unexplored.

During P deficiency, diatoms elevate expression of high affinity phosphate transporters and alkaline phosphatases, maximising P uptake and scavenging [13–17]. This response is coordinated by the transcriptional regulator Phosphate Starvation Response 1 (PSR1) [18], like in green algae [19] and plants [20]. Trypsin proteases are also necessary for diatom acclimation to P limitation [21, 22]. During P starvation, diatoms utilise internal P stores e.g. polyphosphate (polyP), facilitating maintenance of growth despite external P deprivation [23]. Additionally, diatoms economise intracellular P usage, substituting phospholipids with non-P containing forms [12]. This can spare ∼10-30% of total P demand [24]. P-limited diatoms also conserve P by decreasing cellular RNA content [25]. P limitation may additionally impact other P-dependent processes, such as cell signalling. Protein phosphorylation is the most common form of post-translational modification, but it is unknown whether cells maintain this crucial signalling mechanism during P limitation. Furthermore, the proportion of cellular P allocated to protein phosphorylation, and how this may change in response to P deficiency is not known. Consequently, this potentially dynamic cellular P pool remains a major uncertainty in current resource allocation models [26, 27].

In transitioning from P deficient to P sufficient growth, like other algae, diatoms exhibit a “phosphate overplus” response, accumulating P in quantities exceeding physiological demand [28–30] and storing it as polyP in vacuolar granules [23]. In doing so, diatoms drive biogeochemical cycling of P in the oceans [31]. In eukaryotes, Vacuolar Transporter Chaperone (VTC) proteins catalyse polyP biosynthesis, transferring phosphate from ATP, thereby tightly coupling ATP and polyP dynamics [32, 33]. In diatoms, transient increases in phosphate availability are sensed via a rapid calcium (Ca^2+^) dependent-signalling response [11], which has not been observed in other photosynthetic eukaryotes [34, 35]. This drives enhancements in nitrogen (N) uptake within 4 h of P resupply, highlighting rapid crosstalk between P-signalling and N metabolism [6, 11, 36]. Reciprocal regulation of P sensing and acquisition by N availability has also been demonstrated [37]. Moreover, functional characterisation of a plasma membrane localised Ca^2+^-dependent protein kinase (*PtCDPK2*) in *Phaeodactylum tricornutum* revealed its key role mediating resilience to P starvation, as mutants exhibited increased physiological stress under low P conditions [38]. However, the molecular basis for this phenotype, and the downstream players governing recovery from P limitation and transduction of phosphate-induced Ca^2+^ signals remain poorly understood. Although physiological and transcriptomic responses can occur within hours of resupply [11, 13], the earliest regulatory components initiating these changes are unknown.

Protein phosphorylation can rapidly alter protein activity, localization and/or stability, on timescales substantially faster than changes in transcript or protein abundance [39–42]. Hence, phosphoproteomic analysis offers a powerful approach to identify the first molecular responders to phosphate resupply. Changes in light, iron and N status drive robust shifts in the phosphoproteome of diatoms [43, 44]. Here, to deduce how environmental P levels influence cellular investment in protein phosphorylation, we compare the phosphoproteome of *P. tricornutum* in P replete and deplete conditions. We also determine molecular players governing recovery from P starvation, by examining the phosphoproteomic responses of P-limited cells 10 minutes of phosphate resupply. Finally, by applying a pharmacological inhibitor of P-Ca^2+^ signalling (ruthenium red (RuR) [11]) prior to phosphate resupply, we unravel Ca^2+^-dependent and independent regulatory components of these responses.

## MATERIALS AND METHODS

### Strains and culturing

*P. tricornutum* CCAP1055/1 (Culture Collection of Algae and Protozoa, Scottish Association for Marine Science, UK) was the background strain used in this study. *Ptcdpk2.1* and *Ptcdpk2.4* were generated as described by Meeda et al., 2026 (39). Cultures were maintained in sterile filtered (0.22 µm) seawater (FSW) from L4 station, Plymouth (UK) with f/2 nutrients [45] –silicate (Si) and vitamins unless otherwise stated. Phosphate added was according to the P treatments. Axenic cells were maintained in a 16:8 h light: dark cycle illuminated with 60-80 µE m^-2^ s^-1^ light at 18°C.

### Proteomics and phosphoproteomics workflow

Methods for sample preparation, labelling, phosphopeptide enrichment, nano-LC mass spectrometry analysis and data processing are given in the Supplementary Materials and Methods.

### Genome mining, phylogenetic analysis and protein localisation prediction

Protein sequences for plant and green algal phosphate transporters were obtained from Ruili et al., (2020). In addition, the DiatOmicBase [47], Diatoms PLAZA v1.0 [48] and *Tara Oceans* Gene Atlas (v2) [49] portals were mined for diatom and protist sequences. For reconstruction of the phylogenetic tree of phosphate transporters, a sequence alignment was made using MAFFT v7 webserver, using default parameters [50] and trimmed manually using MEGAX software [51]. The best-fitting evolutionary model was also identified using MEGAX via corrected Akaike Information Criterion. A maximum likelihood tree was generated using the WAG with freqs (+F) model and 4 threads with 100 non-parametric bootstrap replicates, with a final alignment of 464 amino acids. For the extended tree, the same method was employed except the final alignment was 436 amino acids, and a LG (+Freq) model was used [52]. Protein localisation predictions were carried out using HECTAR, ASAFIND 2.0, Deeploc2.1 and TargetP 2.0, using default parameters [53–56].

### Localisation of Pt39515 and PtCDPK3 in *P. tricornutum*

Localisation constructs were generated by a modified method from Murphy et al. (2026), as outlined in detail in the Supplementary Materials and Methods, alongside staining procedures using DAPI and the vacuolar membrane dye MDY-64.

### Experiments quantifying ATP and polyP levels of *P. tricornutum* cells under different P regimes

ATP quantification was carried out using a PhosphoWorks™ Fluorimetric ATP Assay Kit (catalog no. 21620, AAT Bioquest, USA), following the manufacturer’s protocol using an ATP standard curve for calibration. Full details of the experimental design and sampling are given in the Supplementary Materials and Methods. Methods for polyP extraction and quantification using the JC-D7 stain are also provided.

## RESULTS

### Overview of the phosphoproteomics experiment

To investigate proteomic and phosphoproteomic changes in *P. tricornutum* in response to P limitation and subsequent replenishment, cells were grown for four days in high (36 µM), low (1.8 µM) or phosphate resupply treatments (i.e. low grown cells harvested 10 minutes after resupply with 36 µM phosphate) (**Figure S1)**. Additional treatments applying RuR prior to resupply, and a low phosphate +RuR control, were also included to elucidate Ca^2+^-dependent components of the phosphate resupply response. Overall, 5415 proteins across the five conditions were identified (satisfying a protein FDR confidence of <5%). A total of 835 and 1497 proteins showed significantly (*p*<0.05) decreased and increased abundance respectively, in low versus high phosphate treatments (**Figure 1A-B; Data S1**). Alterations in the proteome included signature responses to P starvation (**Figure 1C**). As previously reported, N uptake and assimilation proteins also showed significantly decreased abundance [11] (**Figure 1A and D**). Furthermore, proteins involved in Ca^2+^ influx, efflux and decoding were detected (**Figure 1E**). This included identification of a greater number of Ca^2+^ channels (12 in total) than previous proteomics work [11]. Of these, 7 showed significantly increased abundance under low P, alongside 6 CDPKs, and a calmodulin (CAM1), with the Ca^2+^ efflux transporter ECA1 significantly decreased. Together, these results demonstrate our set-up successfully yielded a high-quality proteomics dataset demonstrating characteristic P-starvation responses to low P conditions.

**Figure 1.**
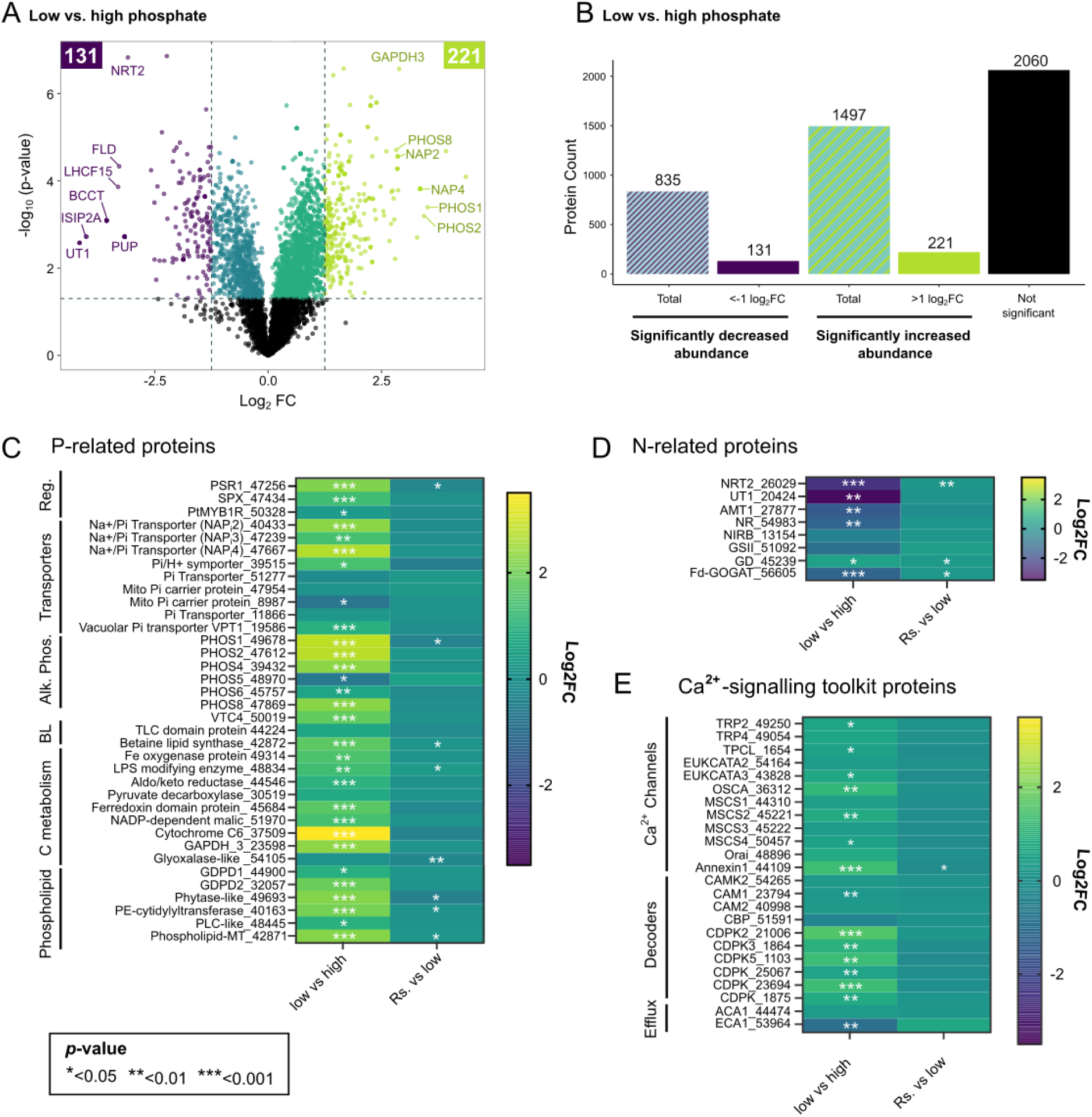
Differential protein abundance in *Phaeodactylum tricornutum* grown in low versus high phosphate conditions and 10 minutes after phosphate resupply. **A**. Volcano plot of proteins showing increased and decreased abundance in *P. tricornutum* in low (1.8 µM) versus high (36 µM) phosphate conditions. The X-axis is the log2fold change (log2FC) (low phosphate versus high phosphate) and Y-axis is the negative log10 of the p-value for paired t-tests between the conditions described (*n*=3). Proteins showing no significant change in abundance (p>0.05) are in black. Proteins with functional annotations showing the greatest fold changes in abundance are given, including: nitrate transporter (NRT2), urea transporter (UT1), Betaine/Carnitine/Choline Transporter (BCCT), iron starvation protein 2A (ISIP2A), flavodoxin (FLD), light harvesting complex protein (LHCF15), purine permease (PUP), alkaline phosphatases (PHOS1, PHOS2 and PHOS8), Na+/phosphate-transporter (NAPi2 and NAPi4) and Glyceraldehyde-3-phosphate dehydrogenase (GAPDH3). **B**. Summary of proteins detected across low and high phosphate treatments grouped by significance and fold change. The total number of proteins with no significantly altered abundance, versus those significantly increased or decreased in abundance (p<0.05), including the number above 1Log2FC or below -1Log2FC are indicated. **C-E**) Heatmaps of relative protein abundance comparing low versus high conditions, as well as resupply (Rs) versus low cells plotting P-related (C) (as annotated in Cruz de Carvalho et al., 2016 [13]), N-related (D), and Ca^2+^-signalling toolkit (E) proteins. The scale bar represents log2FC of protein abundance between the two treatment groups. The p-values (p<0.05: *, p<0.01; ** and p<0.001; ***) for paired t-tests between conditions are also given (*n*=3). Abbreviations and protein names are as follows: phosphate starvation 1 (PSR1), vacuolar transport chaperone (VTC), lipopolysaccaride (LPS), glyceraldehyde-3-phosphate dehydrogenase (GAPDH), glycerophosphoryl diester phosphodiesterase (GDPD), phosphoethanolamine (PE), phospholipase C (PLC), methyltransferase (MT), nitrate transporter (NRT2), urea transporter (UT1), ammonium transporter (AMT1), nitrate reductase (NR), NAD(P)H-dependent nitrite reductase (NIRB), chloroplast-targeted glutamine synthetase, (GSII) (chloroplast-targeted glutamate synthase (Fd-GOGAT), glutamate dehydrogenase (GD), betaine lipids (BL), transient receptor Potential (TRP), two pore cation (TPC), mechanosensitive (MSCS), Ca^2+^-dependent protein kinase (CDPK), calmodulin (CAM), Ca^2+^-binding protein (CBP), P2B Ca^2+^-ATPase (ACA1), and P2A Ca^2+^-ATPase (ECA1).

We did not expect to see substantial changes in the proteome within 10 minutes of phosphate resupply to low P grown cells. Just one protein exhibited significantly increased abundance >1log_2_ fold change, and another significantly decreased <-1log_2_ fold change. Both are uncharacterised proteins based on Uniprot (**Figure S2**). However, strikingly, several proteins associated with P and N metabolism alongside Ca^2+^ signalling, did show significantly altered abundance (*p*-value<0.05) following phosphate resupply even though their log_2_ fold change did not meet the criteria of less than -1 or greater than 1. This included significantly decreased abundance of PSR1, alkaline phosphatase PHOS1, as well as glyoxalase-like and phytase-like proteins (**Figure 1C**). Abundance of two proteins associated with phospholipid biosynthesis, betaine lipid synthase, alongside an LPS-modifying enzyme, significantly increased. Notably, this was in addition to three N-assimilation proteins (**Figure 1D**). Finally, abundance of the Ca^2+^ channel annexin (ANN1) significantly decreased (**Figure 1E**). Hence, a cohort of proteins relating to P metabolism, Ca^2+^-signalling and N uptake/assimilation show rapid significant modifications in abundance within just 10 minutes of phosphate resupply.

### *P. tricornutum* economises P usage by widespread protein dephosphorylation under P deprivation

Of the 5415 proteins detected with corresponding abundance data, 3088 phosphosites were identified that spanned 1122 proteins. Over 500 of the phosphorylated proteins had only one detectable phosphosite, and the number of proteins identified with phosphosites decreased as the number of phosphosites per protein increased (**Figure S3A-B**). The vast majority of phosphosites were on a serine (S) followed by threonine (T) and tyrosine (Y) residues, albeit it was not possible to determine over 500 phosphosites by amino acid (**Figure S3C**). To analyse how phosphorylation of individual phosphosites varied by P regime we normalised the abundance of detected phosphopeptides to respective protein abundance data for each condition, enabling us to discern relative changes in phosphorylation status. Using this approach, a positive adjusted phosphopeptide value indicates that, when considering changes in protein abundance, there is an overall enhancement in the proportion of the protein that is phosphorylated at that phosphosite. In general, we observed substantial and widespread reductions in protein phosphorylation in low versus high phosphate treatments (**Figure 2A**). This included 1346 phosphosites showing significantly (*p*<0.05) reduced phosphorylation (521 of which had a <-1Log_2_ fold change). By comparison, 151 phosphosites had significantly increased phosphorylation (**Figure 2B; Data S2**), just 37 of which showed a greater than 1 Log_2_fold change (p<0.05). After 10 minutes of phosphate resupply, the trend was reversed (**Figure 2C**). More than 660 phosphosites exhibited significantly increased phosphorylation in resupply versus low P cells, 153 of which were above a log_2_FC of 1 (*p*<0.05) (**Figure 2D**). In contrast phosphosites exhibiting decreased phosphorylation were far fewer, with just 10 identified showing a significant log_2_ fold change below –1. These data indicate that P regime drives rapid and substantial modifications in the phosphorylation status of many proteins in *P. tricornutum*. Of the 163 phosphosites showing significantly altered phosphorylation in response to resupply (including those showing a <-1Log_2_ or >1 fold change), 34 were significantly affected by RuR application (i.e. resupply +RuR versus resupply -RuR) (**Dataset S3**). However, to account for non-specific effects of RuR on the phosphoproteome of low grown cells (i.e. without phosphate resupply), we were stringent in discounting 14 phosphosites that also showed significantly altered phosphorylation in the low phosphate +RuR versus low phosphate -RuR control treatment. Hence, pharmacological inhibition of P-Ca^2+^ signalling perturbed P-resupply driven changes in the phosphorylation status of a cohort of around 20 phosphosites, suggesting Ca^2+^-dependent regulation.

**Figure 2.**
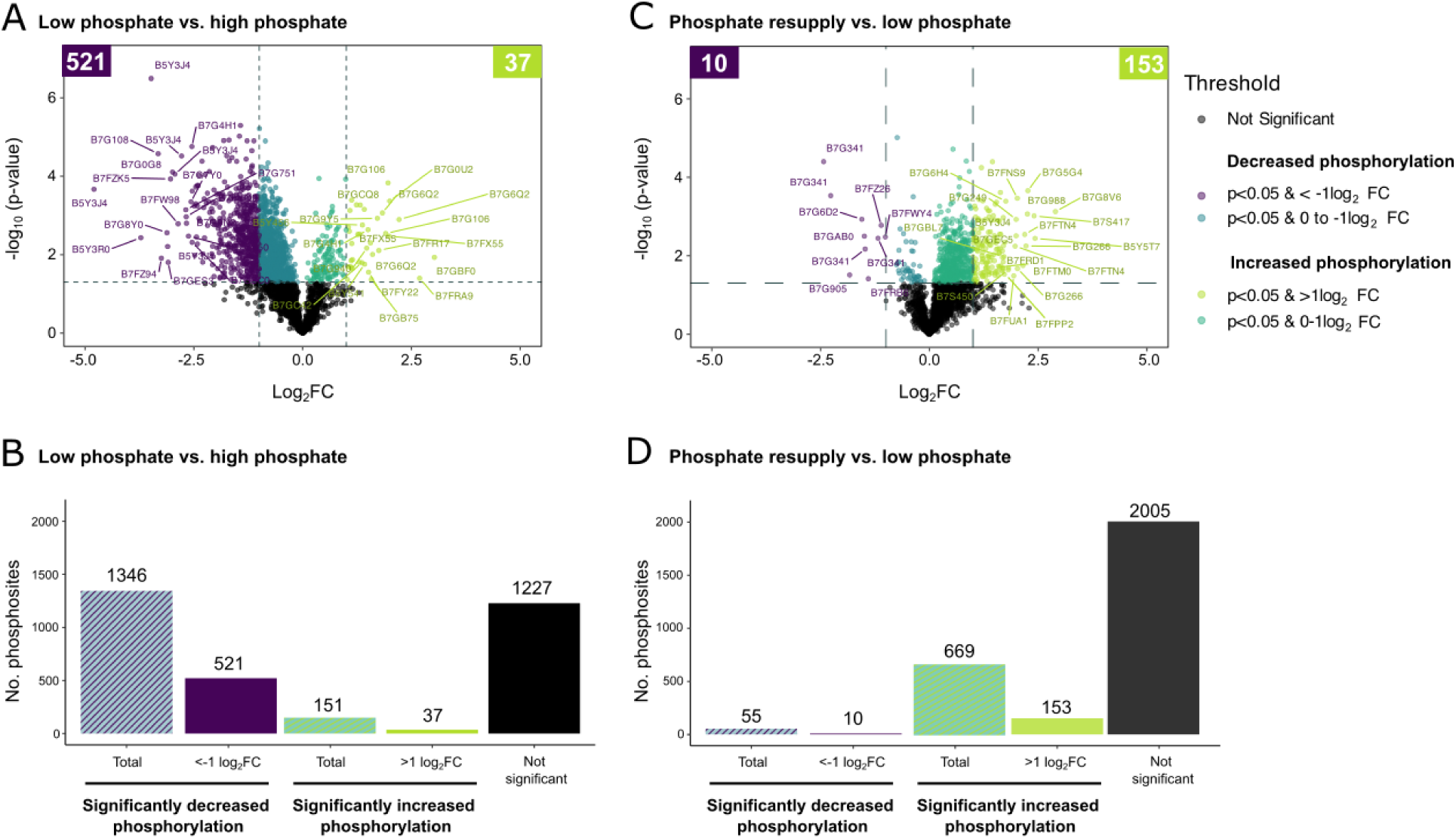
Dynamic shifts in *Phaeodactylum tricornutum* proteome phosphorylation driven by P availability. **A.** Volcano plot showing changes in phosphorylation of phosphopeptides detected in low (1.8 µM) versus high (36 µM) phosphate treatments, plotting -log10 *p*-value against the log2 normalised phosphopeptide abundance (calculated as detailed in the Methods) (*n*=3), with statistical significance determine using paired t-tests between the conditions described. Phosphopeptides showing no significantly altered phosphorylation are represented in black. Uniprot accession numbers are given for top phosphosites showing the greatest fold changes. **B**. Number of phosphopeptides quantified in low versus high P treatments, grouped by significance and log2 foldchange. **C**. Volcano plot as in A but showing phosphopeptides detected in P resupply compared to low P grown cells. **D**. Number of phosphopeptides quantified in resupply versus low P treatments, grouped by significance and log2 foldchange (log2FC).

### Differential phosphorylation of nutrient acquisition and signalling proteins

We conducted a targeted analysis to deduce whether changes in phosphorylation observed in low versus high phosphate treatments affected specific proteins involved in P and N metabolism, and Ca^2+^ signalling. Multiple phosphosites were frequently identified within individual proteins (**Figure 3**). Specific phosphosites showing reduced phosphorylation in low versus high P conditions represented proteins associated with P metabolism, including the plasma membrane phosphate transporters NAP_i_2 (Amb_572-585), SPX protein (S57, S344, S357a, S360; S357 and Amb_350-361), VTC4 (all four detected phosphosites), as well as PE-cytidylyltransferase (S156) for phospholipid remodelling (**Figure 3A**). A phosphosite within the plasma membrane urea transporter (UT1, S688a) [57] (**Figure 3B**) as well as several Ca^2+^-toolkit proteins also showed decreased phosphorylation under P deprivation, including multiple sites within three mechanosensitive (MSCS) channels, two pore cation like (TPCL) (S9 and T1033), ANN1 (S53), in addition to several sites within two Ca^2+^-dependent protein kinases (CDPK3 and CDPK_25067) (**Figure 3C**).

**Figure 3.**
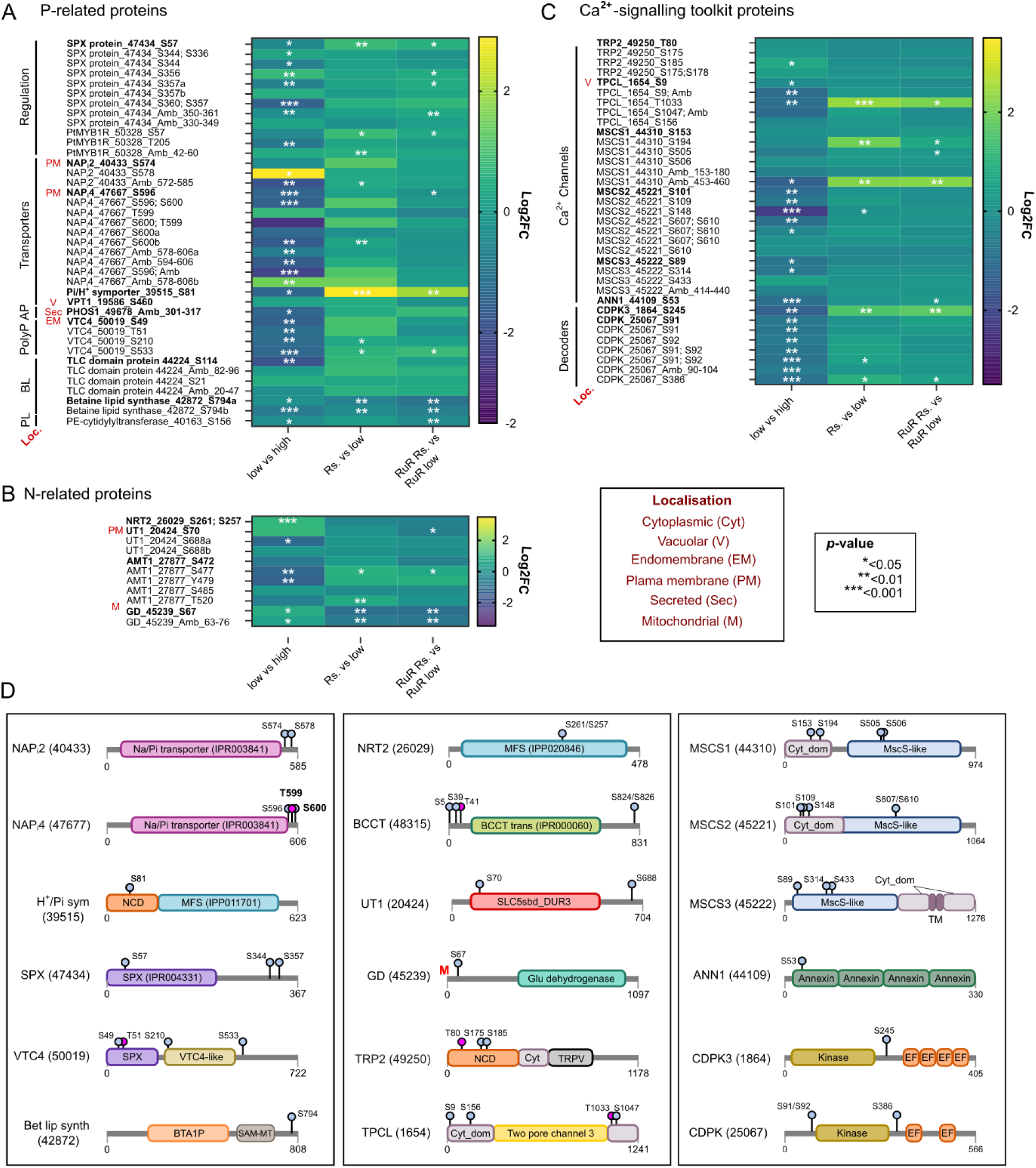
Differential phosphorylation of proteins associated with N and P metabolism and signalling governed by P regime. Heatmaps comparing relative phosphorylation of phosphosites in low (1.8 µM) versus high (36 µM) phosphate conditions, as well as resupply (Rs) versus low cells. An additional comparison of resupply cells pre-treated with ruthenium red (RuR) versus RuR treated low cells is also shown. Log2fold change (Log2FC) normalised phosphopeptide abundance between conditions described, with statistical significance determined using paired t-tests (*n*=3) between the conditions indicated (*p*<0.05: *, *p*<0.01; ** and *p*<0.001; ***). Targeted analyses of proteins involved in P and N metabolism (**A** and **B,** respectively) and Ca^2+^-signalling toolkit (**C**) proteins are shown. The scale bar represents Log2FC of phosphorylation status between the treatment groups. Individual position of phosphosites on each protein are provided, except where the precise phosphosite location is ambiguous (Amb), in which case the region of the peptide within which the phosphosite was identified is given. In some instances, it was not possible to resolve the exact phosphosite location, and in these cases the region of the peptide within which the phosphosite was identified is given. Protein localisation information is given when available from published experimental studies: vacuolar (V), plasma membrane (PM) endomembrane (EM), mitochondrial (M), secretory (Sec) [38, 59–61]. Abbreviations and protein names are as follows: vacuolar transport chaperone (VTC), phosphoethanolamine (PE), nitrate transporter (NRT2), urea transporter (UT1), ammonium transporter (AMT1), glutamate dehydrogenase (GD), and betaine lipids (BL), transient receptor Potential (TRP) channel, two pore cation (TPC)-like channel, mechanosensitive (MscS) channel, Ca^2+^-dependent protein kinase (CDPK), calmodulin (CaM), Ca^2+^-binding protein (CBP). **D.** Protein domain structure as deduced through Interpro analysis [62], with details of phosphosite location (where fully resolved) are indicated. Abbreviations: non-cytoplasmic domain (NCD), major facilitator superfamily (MFS), SYG1, Pho81 and XPR1 (SPX) domain, S-adenosyl-L-methionine-dependent methyltransferases (SAM MT), Betaine/Carnitine/Choline Transporter (BCCT) Family, EF-hand (EF). Red ‘M’ indicates a mitochondrial signal peptide was predicted [55].

However, often phosphosites within a given protein showed opposing phosphorylation patterns. For instance, whereas phosphosite Amb_572-585 in the Na^+^/phosphate transporter, NAP_i_2, was strongly and significantly dephosphorylated under low P conditions, a second phosphosite, within a similar vicinity (S578), showed the greatest increase in phosphorylation of all phosphosites detected under low P conditions **(Figure 3A; Dataset S2**). Strikingly, several other phosphosites exhibited increased phosphorylation, despite cellular P deprivation, suggesting crucial roles during P limitation. This included in NAP_i_4 (Amb_578-606b) and SPX protein (S356 and S344; S336) and notably N assimilation/transport machinery NRT2 (S261; S257) and glutamate dehydrogenase (S67, Amb_63-76) (**Figure 3B**). Just one Ca^2+^-toolkit protein showed increased phosphorylation, TRP2 (S185) (**Figure 3C**). Combining Interpro and PROTTER analyses [58] the location of phosphosites was examined within the context of structural and functional features. In NAP_i_2 and NAP_i_4, all the phosphosites clustered in C-terminus regions, beyond the Na^+^/phosphate transport domain (**Figure 3D)** in a predicted cytoplasmic region (**Figure S4**). Phosphosites within predicted cytoplasmic regions of membrane bound proteins were similarly detected for NRT2 and UT1 (**Figure S4**), alongside Ca^2+^ channels MSCS1-3 (**Figure S5**). Notably, all five phosphosites for AMT1, were located in the C terminus intracellular domain (**Figure S4**). By contrast, the predicted extracellular domain of TRP2 contained 3 phosphosites. In several instances phosphosites were in key functional domains e.g. the SPX domain of both SPX protein and VTC4, or the major facilitator superfamily (MFS) domain of NRT2 (**Figure 3D)**.

### A novel intracellular phosphate-H^+^ symporter exhibits the greatest increase in phosphorylation observed following P resupply

Of the 723 phosphopeptides showing significantly altered phosphorylation in response to phosphate resupply, the majority (669) exhibited significantly increased phosphorylation (**Figure 2D**). This included phosphosites in many proteins associated with P metabolism, such as within the SPX domain (S57) of SPX protein, as well as sites within VTC4 (S210, S533), phosphate transporters NAP_i_2 (Amb_572-585), NAP_i_4 (S600b) and predicted H^+^/phosphate transporter (Pt39515_S81), alongside MYB1R _50238 (S57; Amb_42-60) (**Figure 3A**) that is closely related to PtPSR1 [18]. This was also the case for key N acquisition (e.g. AMT1_S477) and Ca^2+^-signalling tool-kit proteins (**Figure 3B-C**). These changes, that occurred within 10 minutes of P resupply, suggest targeted phospho-regulation of P and N acquisition and signalling machinery as some of the very first responses to P replenishment.

The phosphosite exhibiting the greatest increase in phosphorylation following P resupply was S81 of the predicted H^+^/phosphate symporter (Pt39515) (**Figure 3A**; **Figure 4A-B**). This protein also shows significantly increased abundance in low versus high P conditions (**Figure 1A**; **Figure 4C**), and while there is no significant change in the abundance of this protein following phosphate resupply, there was a 2.9 log_2_fold increase in its phosphorylation status (**Figure 4B-C**). Phylogenetic analysis indicates that Pt39515 is distinct from *P. tricornutum* Na^+^/phosphate co-transporters: NAP_i_1-NAP_i_6 and is instead more closely related to transporters from the plant PHT1 /green algal PTA clade [63, 64] (**Figure 4D**). *P. tricornutum* possesses two additional PHT1 homologues (Pt46251 and Pt17265 (Hp_i_1)) (blue clade, **Figure S6A**), alongside two closely related proteins including Pt39515 and Pt46692 that cluster in distinct groupings we have termed PHT1-A and PHT1-B, respectively (**Figure S6A**). Pt39515 is strongly conserved, with orthologues present in diverse diatom genomes, including environmentally prevalent *Tara Oceans* Metagenome and Single Cell Assembled Genomes (SMAGs) (**Figure S6-S7**). Whilst forming a distinct clade, Pt39515 orthologues have retained conserved residues necessary for phosphate binding [65, 66] (**Figure S6B**), suggesting that they are phosphate transporters. PHT1-A clade sequences are predicted to be endomembrane localized, either in the lysosome/vacuole, Golgi, or ER (**Data S4**) [53–56]. To further investigate this, we generated transgenic cells expressing Pt39515 fused to a C-terminal mApple tag, under the control of its endogenous promoter region. This revealed Pt39515-mApple fluorescence in a punctate region in the vicinity of the chloroplast in low-phosphate grown cells (**Figure 4E**), suggesting localisation to a membrane compartment associated with the plastid. Faint signal at the cell periphery was also detectable in low-phosphate grown cells. Quantifying average mApple fluorescence per cell, confirmed enhanced expression in low versus high phosphate conditions, exceeding background fluorescence detected in WT (**Figure 4F; Figure S8)**, and supporting the proteomics (**Figure 1A**; **Figure 4C**). Taken together, our analysis identifies that phospho-regulation of a conserved novel intracellular H^+^/phosphate transporter is one of the first detectable steps following P resupply, and likely plays an important role in internal phosphate trafficking in P-starved diatom cells upon P replenishment.

**Figure 4.**
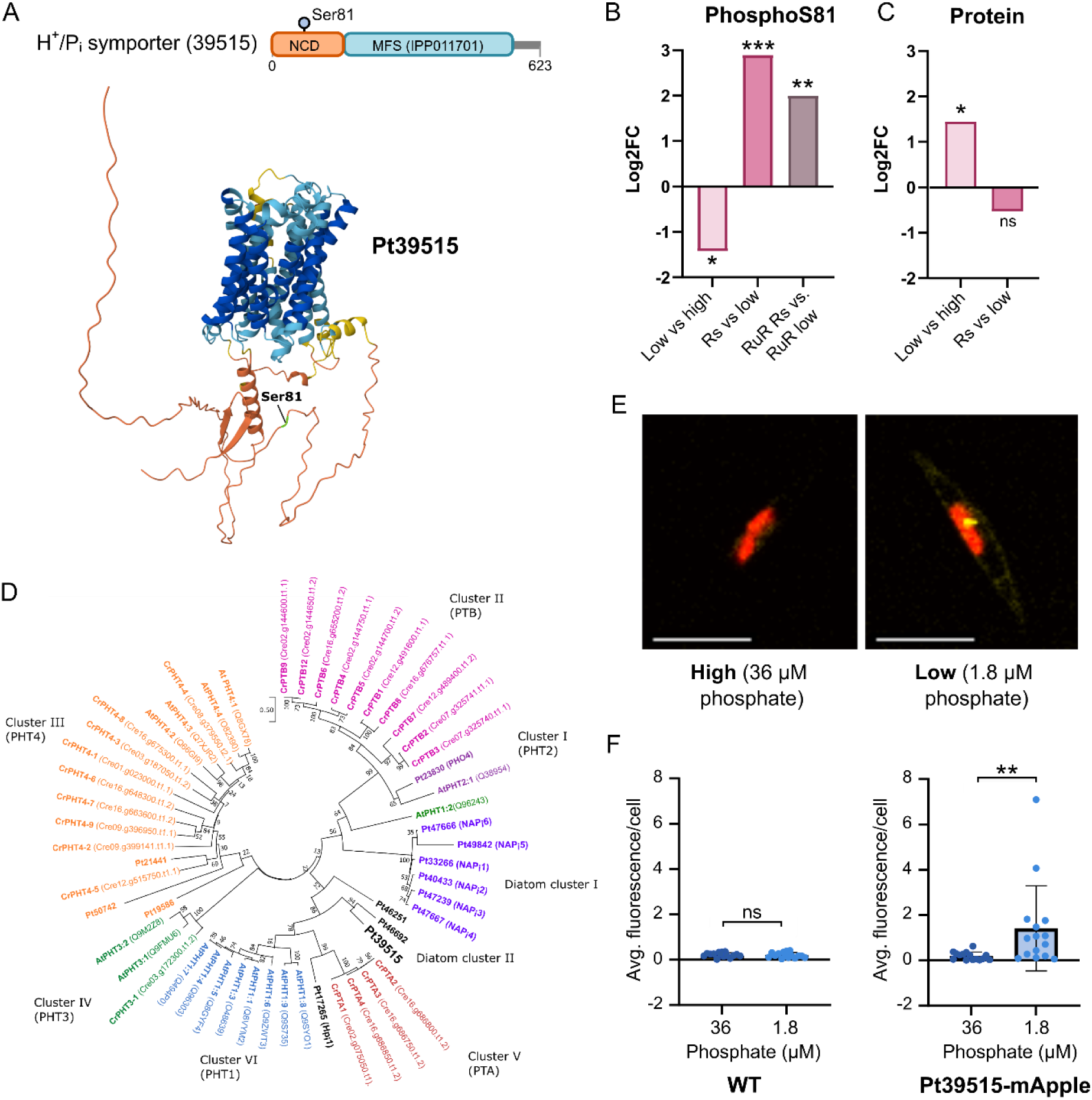
A novel intracellular phosphate-H^+^ symporter shows substantially altered phosphorylation following phosphate resupply. **A.** Protein domain structure of phosphate-H^+^ symporter Pt39515 with with phosphosite S81 indicated. An Alphafold3 model of Pt39515 indicating the position of S81 is given below. The confidence level of the prediction is indicated by colour, as follows: dark blue (very high, pLDDT > 90), confident (90 > pLDDT > 70), low (70 > pLDDT > 50) and very low (pLDDT < 50), where pLDDT is a per-atom confidence estimate on a 0-100 scale where a higher value indicates higher confidence. Abbreviations: non-cytoplasmic domain (NCD), major facilitator superfamily (MFS). **B**. Log2fold change (log2FC) in normalised phosphopeptide abundance of Pt39515_S81, with statistical significance determined using paired t-tests (*n*=3) between the treatment groups indicated (*p*<0.05: *, *p*<0.01; ** and *p*<0.001; ***). **C**. Log2FC changes in protein abundance of Pt39515 in low versus high phosphate grown cells and following phosphate resupply (as also plotted in the heatmap in Figure 1C), indicating the *p*-value for paired t-tests between the conditions described (*n*=3) between the conditions indicated (*p*<0.05: *, *p*<0.01; ** and *p*<0.001; ***). **D.** Phylogenetic relationship of *P. tricornutum* (Pt) phosphate transporter proteins compared to green plants and algae. Phylogenetic tree was reconstructed using MEGAX using the maximum likelihood method with 100 bootstraps WAG with freqs (+F) model and 4 threads. The ML heuristic method employed was nearest-neighbour-interchange (NNI). Cr: *Chlamydomonas reinhardtii;* At: *Arabidopsis thaliana.* PHT: phosphate transporter. IDs were taken from Uniprot and *C. reinhardtii* locus number [63]. **E.** Confocal microscopy of Pt39515-mApple expressed under the control of its native promoter, grown with 36 µM (high) or 1.8 µM (low) phosphate conditions for four days. Chlorophyll autofluorescence is red, whereas mApple fluorescence is yellow. Scale bar: 10 µm. **F.** Average mApple fluorescence/cell quantified in wildtype (WT) and Pt39515-mApple cells grown for four days with 36 µM (high) or 1.8 µM (low) phosphate. Data were analysed from *n*=3 biological replicates per treatment per strain, with 5 technical replicate images analysed per biological replicate. Individual data points represent the average fluorescence/ cell of all cells examined in each image (i.e. 15 images in total per strain treatment), with a minimum of 5 cells examined per image, error bars are SDM. Results of a non-parametric two-tailed Mann-Whitney t-test are indicated: non-significant (ns); p<0.01 (**).

### A cohort of mitochondrial proteins show altered phosphorylation following phosphate resupply

We identified multiple predicted mitochondrial proteins among the phosphosites showing the greatest changes in phosphorylation following resupply (**Figure 5A**). This included T36, within the mitochondrial ATP-synthase (γ-subunit) that exhibited significantly increased phosphorylation upon P resupply. This was in addition to T41 of a predicted mitochondrial carrier protein (MCP; Pt39648), as well as phosphosites on a ubiquitin carboxyl-terminal hydrolase (Pt42571). The latter contains a predicted mitochondrial signal peptide [55] and shows significantly decreased abundance 10 minutes following P resupply (**Figure 5B**). In contrast, pyruvate dehydrogenase (PDH, S305), a crucial enzyme for the citric acid cycle in the mitochondrial matrix, becomes significantly dephosphorylated following P resupply (**Figure 5A).** This is also the case for glutamate dehydrogenase (GD), and both enzymes showed significantly altered abundance upon P resupply (**Figure 5B**). Other proteins also exhibited decreases in phosphorylation included 5 phosphosites within an ABC transporter with predicted mitochondrial ATP-binding domain (Pt21548) and S1244 of a predicted mitochondrial metal transporter (MMT1) (**Figure 5C**). Notably, several of these changes in phosphorylation were significantly affected by RuR (**Data S3)**. This included phosphosites within the mitochondrial ATP-synthase (γ subunit) as well as mitochondrial ubiquitin carboxyl-terminal hydrolase (UCH) and glutamate dehydrogenase. This was also the case for phosphoglycerate kinase, the first ATP-generating enzyme of glycolysis in the cytosol. This data suggests that these phosphorylation events exhibit Ca^2+^ dependency. As mitochondria are key sites for ATP biosynthesis, we quantified cellular ATP levels. We did not detect significantly different ATP levels between high and low grown cells (**Figure 5D**). However, we observed rapid and significant decreases in ATP per cell within just 1 h following phosphate resupply to low grown cells and this became even more pronounced within 4 h (**Figure 5E**). These data indicate a net ATP depletion in the first instances of P replenishment to P-starved cells.

**Figure 5.**
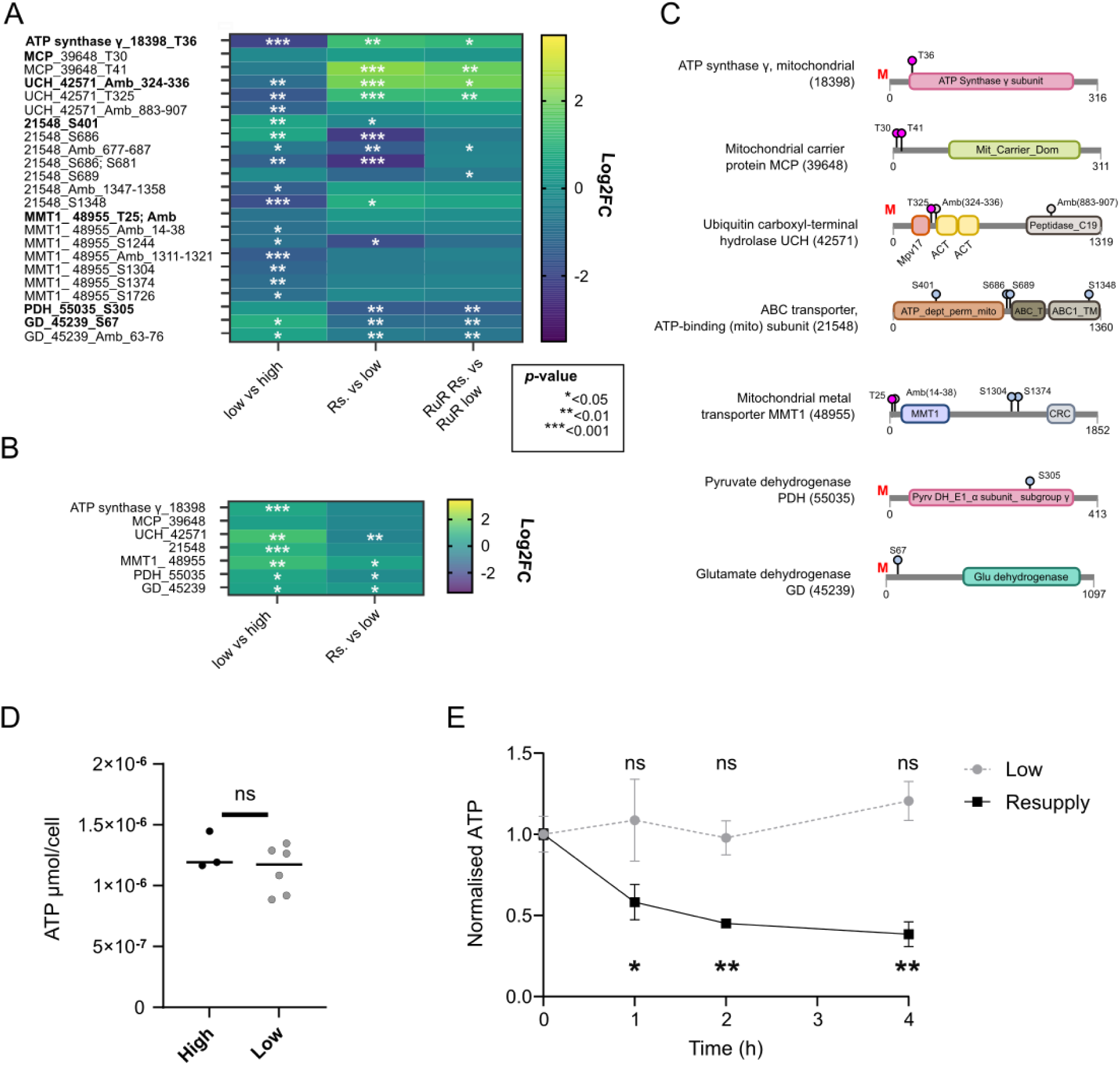
A cohort of mitochondrial proteins show altered phosphorylation following phosphate resupply. **A.** Heatmap comparing relative phosphorylation of phosphosites of mitochondrial associated proteins, comparing low (1.8 µM) versus high (36 µM) phosphate conditions, as well as resupply (Rs) versus low cells. Results comparing resupply +RuR versus low +RuR are also shown. Log2fold changes (FC) indicate log2 normalised phosphopeptide abundance between the treatment groups, with statistical significance determined using paired t-tests (*n*=3) between the conditions indicated (*p*<0.05: *, *p*<0.01; ** and *p*<0.001; ***). Abbreviations: mitochondrial carrier protein (MCP), ubiquitin carboxyl-terminal hydrolase (UCH), mitochondrial metal transporter 1 (MMT1) related, pyruvate dehydrogenase (PDH) and glutamate dehydrogenase (GD). **B.** As in A but comparing changes in relative protein abundance in low P versus high P and resupply (Rs) versus low P cells. The scale bar represents log2 fold change (FC) of protein abundance between the treatment groups. **C.** Protein domain structure of phosphoproteins of interest as deduced through Interpro analysis, with phosphosite location (when unambiguous) indicated. Red ‘M’ indicates a mitochondrial signal peptide was predicted via HECTAR [55]. **D**. Concentration of ATP (µmol/cell) in 6-day old *P. tricornutum* cells grown in high (36 µM) or low (1.8 µM) phosphate conditions. Results of a t-test comparing high versus low grown cells were not significant (ns) (n=3 and 6 for high and low treatments, respectively). **E.** A time-course of cellular ATP levels 1, 2 and 4 hours after phosphate resupply (36 µM) to 6-day old low grown cells. The control treatment (low) without phosphate resupply is also shown. Cellular ATP concentrations were normalised to t=0 for each respective treatment. Results of a two-way repeated-measures mixed-effects ANOVA with Holm-Šidák multiple-comparisons correction comparing each time-point to t=0 for each treatment (not significant, ns; p<0.05: *, p<0.01; n=3) are shown (mean ± SDM).

### Ca^2+^- and CDPK-dependent regulation of luxury polyP dynamics

Finally, we saw significantly increased phosphorylation of multiple Ca^2+^-toolkit proteins following phosphate resupply (**Figure 3C**). This included three phosphosites within two CDPKs (PtCDPK3 and PtCDPK_25067). Both S245 (PtCDPK3) and S386 (PtCDPK_25067) are located between the predicted Ser/Thr kinase domain and the EF hands, within the autoinhibitory junction [67] (**Figure 3D**). PtCDPK3 showed significantly increased protein abundance and was significantly dephosphorylated at S245 under low versus high phosphate conditions (**Figure 6A)**. S245 then became rapidly re-phosphorylated upon P resupply, even though protein abundance did not change. To further investigate the role of PtCDPK3, we generated transgenic cells expressing PtCDPK3 fused to mApple, under the control of its endogenous promoter region. Strikingly, PtCDPK3-mApple fluorescence was localised to the vacuolar membrane (**Figure 6B**), confirmed via co-staining with the tonoplast stain, MDY-64 (**Figure S9**). Consistent with this, we identified a dileucine-based tonoplast targeting motif at the protein N terminus [68]. mApple fluorescence per cell was significantly higher in the low P treatment (**Figure 6C**), supporting the proteomics data (**Figure 1E**; **Figure 6A**). Increased fluorescence corresponded with the frequent appearance of bright rings within the vacuoles (**Figure 6B; Figure S10**) that strongly diminished within 1-2 hours of phosphate resupply (**Figure 6D; Video S1**). Diatoms accumulate polyP in large apical vacuolar granules [23]. We therefore examined the localisation of PtCDPK3-mApple with respect to intracellular polyP stores. Notably, in the high phosphate treatment, polyP was detected in punctate regions associated with PtCDPK3-mApple spots (**Figure 6E**). PolyP was less evident in the 1.8 µM phosphate treatment, but became visible again 4 h after phosphate resupply i.e. coinciding with declining ATP levels following P resupply (**Figure 5E**). PtCDPK3-mApple signal strongly overlapped with the DAPI-polyP (**Figure S11**). Quantification using the polyP-specific stain JC-D7 [69, 70], detected significant increases in polyP content per cell within just 30 mins of phosphate resupply, with levels plateauing after 2-4 h (**Figure 6F**, inset). To test whether P-Ca^2+^ signalling regulates this response, we examined the impact of RuR on polyP accumulation. Rapid recovery of polyP levels within 2 h of phosphate resupply was significantly inhibited by 88% in the +RuR versus -RuR treatment (**Figure 6F**). This strongly suggests that the response is Ca^2+^-dependent and regulated by the P-Ca^2+^ signalling pathway.

**Figure 6.**
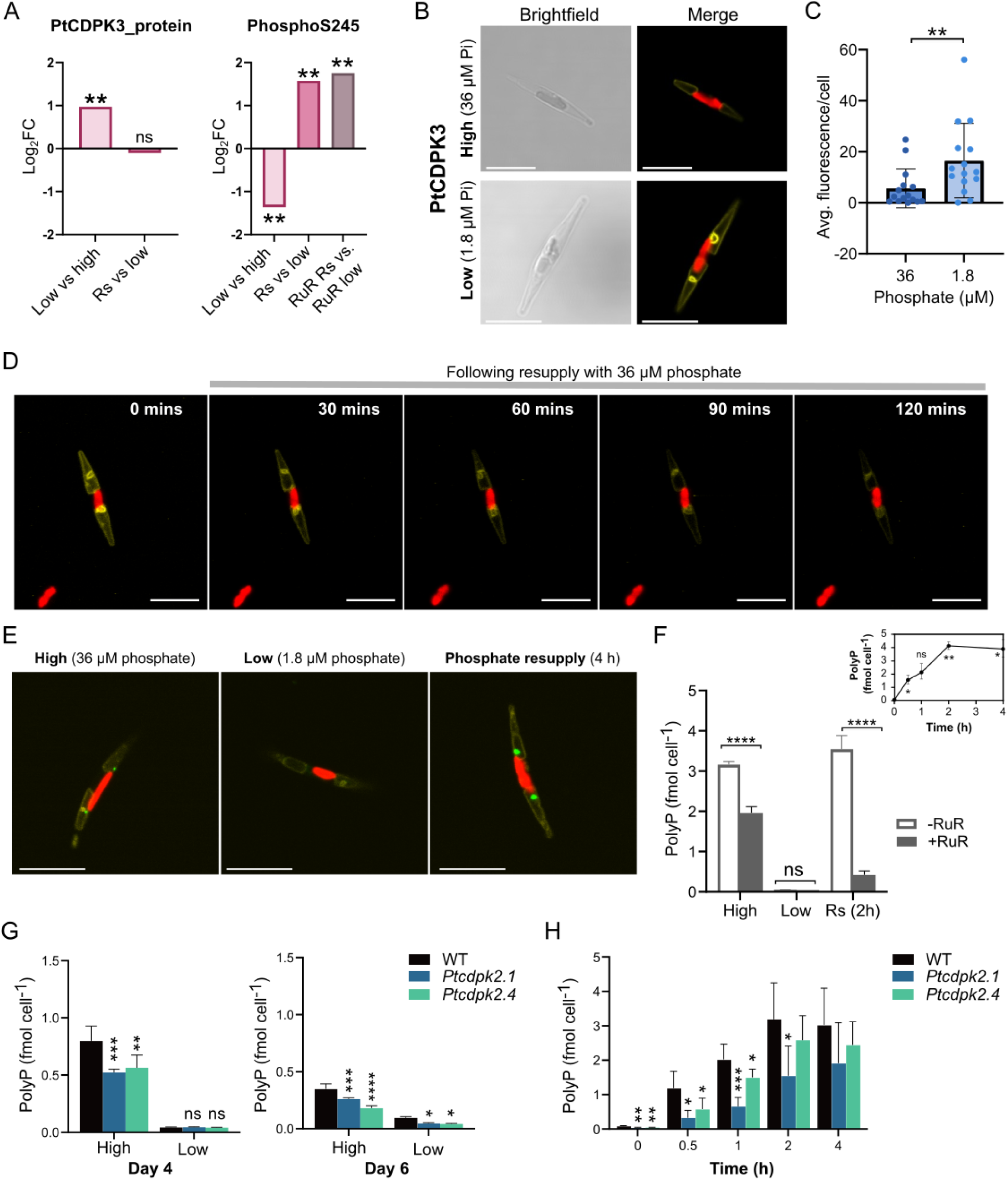
A role for Ca^2+^- and CDPK-signalling in regulating vacuolar polyP dynamics. **A**. Log2FC changes in protein abundance of PtCDPK3 in low versus high phosphate grown cells and following phosphate resupply (as also plotted in the heatmap in Figure 1C), indicating the *p*-value for paired t-tests between the conditions described (*n*=3) between the conditions indicated (*p*<0.05: *, *p*<0.01; ** and *p*<0.001; ***) (right). Log2fold change (log2FC) in normalised phosphopeptide abundance of PtCDPK3_S245, with statistical significance determined using paired t-tests (*n* = 3) between the treatment groups indicated (*p*<0.05: *, *p*<0.01; ** and *p*<0.001; ***) (right). **B.** Confocal microscopy of PtCDPK3-mApple expressed under the control of its native promoter, grown with 36 µM (high) or 1.8 µM (low) phosphate conditions for four days. Scale bar: 10 µm. **C.** Average mApple fluorescence/cell quantified in PtCDPK3-mApple cells grown in the conditions described in (A). Data were analysed from *n*=3 biological replicates per treatment per strain, with 5 technical replicate images analysed per biological replicate, error bars are SDM. Individual data points represent the average fluorescence/cell of all cells examined in each image (i.e. 15 images in total per strain treatment), with a minimum of 5 cells examined per image. **D.** Time-lapse image of PtCDPK3-mApple cell grown for 4 days in low (1.8 µM) phosphate conditions, following resupply with 36 µM phosphate. Timestamps indicate the time (minutes) from the beginning of the perfusion experiment; scale bar 10 µm. **E**. Localisation of PtCDPK3-mApple fluorescence (yellow) and polyP DAPI (405 nm/535 – 570 nm excitation/ emission in green) in 4-day old cells grown in 36 µM (high) or 1.8 µM (low) phosphate, or 4 h following phosphate resupply. Chlorophyll autofluorescence is shown in red. Scale bar: 10 µm. **F**. PolyP per cell of *P. tricornutum* grown under different P regimes. Cells were grown in high (36 µM) or low (1.8 µM) phosphate treatments for 6 days, and harvested 2h following 36 µM phosphate resupply to low grown cells (resupply treatment; Rs (2h)), in the absence or presence of RuR. Results of a two-way ANOVA with Tukey’s multiple comparisons correction are shown, comparing the +RuR (10 µM) and -RuR in the different phosphate treatments (ns, not significant; \*\*\*\**P* <0.0001). Inset displays a separate time-course monitoring polyP per cell 0.5, 1, 2 and 4 h post phosphate resupply to 6-day old low phosphate grown cells (mean ± SDM, *n* = 3). Statistical significance was assessed using a one-way repeated measures ANOVA test with Dunnett’s multiple comparisons correction, comparing results at each time-point to t=0 (ns, not significant;\**P* <0.05, \*\**P* <0.01). **G.** PolyP per cell in WT versus *Ptcdpk2.1* and *Ptcdpk2.4* mutants in high versus low phosphate conditions after 4 and 6 days. Results of a two-way ANOVA with Dunnett’s multiple-comparisons correction comparing each *Ptcdpk2* mutant with the WT within each treatment are shown: ns, not significant; \**P* <0.05, \*\**P* <0.01, \*\*\**P* <0.001, \*\*\*\**P* <0.0001 (mean ± SDM, *n* = 3). **H**. A time-course of polyP content per cell following 36 µM phosphate resupply to 6-day old low phosphate (1.8 µM) grown WT and mutant lines. Data were compiled from two independent biological triplicate experiments (*n* = 6 biological replicates; mean ± SD). Statistical significance was assessed using a two-way repeated-measures mixed-effects ANOVA with Holm-Šidák multiple-comparisons correction (*P < 0.05, **P < 0.01, ***P < 0.001).

Given the newly identified role of P-Ca^2+^ signalling in regulating polyP storage, we sought to determine whether this could explain previously observed phenotypes of mutants of other components of the P-Ca^2+^ signalling pathway. PtCDPK2 is a plasma membrane-localised CDPK strongly upregulated during P deprivation, and loss-of-function mutants exhibit enhanced physiological stress under P starvation [38]. To investigate whether altered polyP homeostasis may contribute to this phenotype, we examined polyP content under the different P regimes. In high phosphate, independent mutants *Ptcdpk2.1* and *Ptcdpk2.4,* exhibited significantly reduced polyP levels relative to WT, with decreases of 34% and 45% respectively on day 4 and 25% and 48% on day 6 (**Figure 6G**). This was despite no detectable differences in growth rate between the strains in these conditions [38]. Reduced levels were also apparent in the low phosphate treatment on day 6. Notably, mutants also showed a significantly diminished capacity to restore polyP compared to WT 0.5-1 h post resupply (**Figure 6H**). These data further support the role of Ca^2+^- and CDPK-dependent signalling in regulating vacuolar polyP dynamics, including immediately following P resupply.

## DISCUSSION

To cope with fluctuating P supply, diatoms employ a sophisticated molecular toolkit for P sensing, storage, and acquisition. However, many players remain unknown. Using global phosphoproteomics, we identify rapid widespread changes in protein phosphorylation in response to P availability. While diatoms remodel phospholipids to economise P under deficiency [12, 24, 25, 71], our study reveals widespread proteome dephosphorylation as a previously unrecognized P-sparing strategy. Proteins constitute ∼30-65% of cell mass and are well known for their high N and carbon content [71, 72]. In contrast, P allocation to the phosphoproteome via phosphorylation is poorly constrained [26, 27]. This is despite proteome-wide studies reporting ∼0.8 mol P/mol protein in other eukaryotes [73], highlighting the phosphoproteome as a potentially substantial P reservoir. Across 5,415 proteins, we identified 3,088 phosphosites (∼1.75 per protein). Assuming ∼100,000,000 protein copies per cell [74], if 44% of phosphosites show reduced phosphorylation under moderate P limitation (i.e. grown with 1.8 µM phosphate) whereas only 4.9% exhibit increased phosphorylation, we estimate that this could redistribute up to 0.114 fmol of P. Although this assumes equal protein abundances and complete occupancy under P-replete conditions of the phosphosites showing reduced phosphorylation under P limitation, it nevertheless indicates that protein phosphorylation may represent a non-trivial, dynamic component of cellular P allocation. Notably, this could correspond to up to ∼5.7% of the total P quota of an 8 µM phosphate grown *P. tricornutum* cell (2 fmol P/cell) [75]. Consequently, whilst nucleic acids, phospholipids and polyP undoubtedly account for most cellular P content [26, 71], redistribution of phosphoproteomic P reserves provides an additional mechanism of flexible P allocation that could yield a competitive advantage under severe P limitation. In doing so, P scarcity directly shapes cell signalling networks, affecting phosphoregulatory control of multiple pathways, including nutrient acquisition and signalling (**Figure S12A**).

Upon P replenishment, substantial phosphoproteome remodelling occurred within minutes. This illuminates coordinated responses to phosphate resupply unfolding on timescales more rapid than captured previously sampling hours or days after P resupply [11, 13]. For instance, significant changes in the abundance of key P acquisition proteins (e.g. PtNAPi2/PtNAPi4) were not observed after 4 h of resupply [11], even though the current study reveals their rapid phospho-regulation within minutes. Moreover, our phosphoproteomic analyses uncover novel components of the P resupply response, including phospho-regulation of proteins for intracellular P transport, mitochondrial and ATP metabolism, alongside a tonoplast localised Ca²⁺-sensor protein (PtCDPK3) associated with vacuolar polyP stores (**Figure S12B**). In parallel, we found cellular ATP declined 1-4 h following P resupply, likely reflecting increased ATP demand for phosphorylation and polyP synthesis that exceeds ATP production. Hence, recovery from P starvation entails rapid regulation of organellar energy metabolism and the P overplus response, to kick-start growth and exploit newly available P.

The phosphosite exhibiting the greatest increase in phosphorylation following phosphate resupply, belongs to a intracellular H^+^/phosphate symporter. This protein, localised to a punctate region near the chloroplast in *P. tricornutum*, is strongly conserved in diverse diatoms. Our evidence thus indicates that intracellular P trafficking and use are key processes rapidly regulated by diatoms upon P replenishment. This was supported by evidence that a range of mitochondrial proteins also exhibited rapid phospho-regulation. Among the rare phosphosites dephosphorylated following P resupply was S305 within pyruvate dehydrogenase (PDH) E1 subunit. PDH complex catalyses the conversion of pyruvate to acetyl-CoA, linking glycolysis, the citric acid cycle and oxidative phosphorylation. In other eukaryotes, Ca^2+^ is an important metabolic regulator of citric acid cycle enzymes [76–79]. Increasing Ca^2+^ levels can enhance PDH phosphatase activity, causing dephosphorylation and thereby activation of PDH complex [80–82]. The S305 within *P. tricornutum* PDH E1 dephosphorylated upon P resupply, corresponds to the conserved phosphorylation site of human PDH E1α (HGH**S**MSDPGV), responsible for activation by dephosphorylation [81, 83]. This drives generation of ATP, as well as NADH, vital for further ATP production through oxidative phosphorylation (**Figure S12B**). Notably, changes in the phosphorylation of Pt39515 and the mitochondrial ATP synthase γ-subunit alongside other mitochondrial enzymes following phosphate resupply were significantly perturbed by RuR, suggesting that phosphate-resupply induced Ca^2+^ signals [11] control their regulation. Ca^2+^-dependent control of ATP synthase γ-subunit and ATP production has previously been reported in animals [84, 85]. Hence, collectively, our results suggest that phosphate, and Ca^2+^, are crucial signals regulating ATP metabolism to restore metabolic activity following P replenishment.

Notably, RuR also caused a substantial inhibition (by 88%) of polyP biosynthesis following P resupply. As ATP is a crucial substrate for polyP production [32, 33], it is possible that this effect is in part governed by inhibition of upstream Ca^2+^-dependent aspects of ATP metabolism. However, identification of the vacuolar membrane associated PtCDPK3 (**Figure 6**), alongside the impaired ability of *Ptcdpk2* mutants to replenish polyP levels, strongly suggests that Ca^2+^ and CDPK-dependent signalling control specific aspects of vacuolar P transport, polyP biosynthesis and/or regulation. Whilst it is well established that diatoms store polyP in granules within large apical vacuoles [23], many components of diatom polyP metabolism remain to be functionally characterised. First discovered in yeast, and present in other algae including diatoms [28, 30, 32, 68, 86], the VTC complex plays a crucial role governing polyP biosynthesis via subunits VTC1-4 [87, 88]. Of these, we detected only PtVTC4 via phosphoproteomics and identified four phosphosites. All were dephosphorylated under P deprivation, with two phosphorylated upon P resupply. In *P. tricornutum*, PtVTC4 is ER localised independent of P status (when over-expressed using the nitrate reductase promoter), like PtVTC1 [86]. However, analogous to PtCDPK3, PtVTC3 is associated with the vacuolar membrane including in ring-like sub-compartments [68]. In yeast, phosphorylation sites have been identified in VTC2 and VTC3 [89]. Substitution of S190 in VTC3, caused a 75% reduction in polyP synthesis. The functional roles of phosphosites within PtVTC4 remain to be established, as well as whether vacuolar-associated PtCDPK3 directly regulates apparatus for polyP production and/or use. By contrast, the role of the plasma membrane-localised PtCDPK2 is more likely to be involved in relaying the P-Ca^2+^ signal at the cell periphery to downstream targets within. PtCDPK2 is also emerging as a regulator of both P starvation responses, and the switch between P deplete and replete states. This could explain why mutants are not entirely defective in polyP biosynthesis but rather show delayed synthesis upon resupply. Our collective findings highlight that P-Ca^2+^ signalling is a network regulator of multiple aspects of P homeostasis.

Maintenance of intracellular P fluxes relies on a suite of P scavenging and acquisition apparatus[86, 90, 91]. Among the eight alkaline phosphatases identified in *P. tricornutum* (PtPHOS1-PtPHOS8), two are secreted [86, 90, 92]. Extracellular release of PHOS1 occurs in P deplete but not replete conditions, and is thought to be regulated via phosphorylation [59]. Consistent with this, PtPHOS1 exhibited increased phosphorylation under P-limiting conditions. In addition to phospho-regulation of alkaline phosphatases, we also identified altered phosphorylation status of plasma membrane Na^+^/phosphate transporters, PtNAP_i_2 and PtNAP_i_4 [86]. In *A. thaliana*, phosphorylation controls trafficking of PHOSPHATE TRANSPORTER1 (PHT1), from the ER to the plasma membrane under low P, to its endocytosis and routing to lytic vacuoles in high P [93]. As PtNAP_i_2 and PtNAP_i_4 have several phosphosites with opposing phosphorylation patterns under different P regimes, they may too govern transporter trafficking and/or recycling, which requires further investigation.

Accumulating evidence highlights the importance of crosstalk between P and N metabolism, to balance acquisition of these crucial macronutrients [6, 22, 36, 37]. Our study now highlights that this interplay also occurs at the post-translational level. Diatom nitrate uptake is mediated by low-affinity nitrate transporter/peptide transporter family (NPF) proteins (formally NRT1) and high-affinity NRT2 protein families [94]. *P. tricornutum* encodes six NRT2 and two NPF genes. One of the few phosphosites exhibiting increased phosphorylation under low P conditions belonged to PtNRT2_26029. Pt26029 abundance was also significantly decreased during P limitation, increasing following P resupply. In plants, phospho-regulatory control of NRT2 is well documented [95]. *A. thaliana* AtNRT2.1 is phosphorylated at five independent sites, governing transporter activation/deactivation [96, 97] or altered protein-protein interactions [98]. Moreover, a Ca^2+^-dependent protein kinase, AtCPK28, directly phosphorylates the N-terminal S21 to positively regulate nitrate uptake [99]. Whilst the functional impact/s of Pt26029 phosphorylation remains to be elucidated, we speculate indispensable roles given the allocation of P despite P limitation. We also identified P-responsive phosphosites within the plasma membrane localised urea transporter PtUT1_20424 [57] and ammonium transporter, PtAMT1_27877. In *A. thaliana*, increased phosphorylation of S450 in AMT1;1 mediated by CPK32 causes increased ammonium uptake [100]. Intriguingly, phosphorylation of PtAMT1_27877 T520 showed Ca^2+^-dependency. This further supports the role of phosphate-induced Ca^2+^ signalling in regulating N transport in *P. tricornutum* [11], though the specific sensor/relay proteins remain to be determined. Moreover, as UT1 and AMT1 also show altered phosphorylation in response to N status and light in *P. tricornutum* [43], these data highlight the complexity and multi-stressor control of post-translational regulation of nutrient acquisition machinery in diatoms.

## Acknowledgments

This work was supported by Natural Environment Research Council (NERC) Independent Research Fellowship NE/R015449/2 and Biotechnology and Biological Sciences Research Council (BBSRC) New Investigator Grant BB/W006286/1. Yasmin Meeda is also grateful for a jointly funded PhD studentship between the University of Exeter and the Marine Biological Association (MBA), Plymouth, UK. Jiayou GE and Qiong Zhang acknowledge support from the Brian Morton Exchange Fund, administered by the MBA.

## Author contributions

Conceptualization: YM, YT, KEH, GLW and AM; Experiments: YM, YT, KEH, SW, KJH, PAL and JGE; Data analysis: YM, YT, KEH, SW, KJH, PAL and JGE; Writing (original draft): KEH and YM; Writing (review & editing): KEH, YM, GLW, AM. Funding acquisition: KEH, JGE and QZ. All authors approved the manuscript.

## Data, code, and materials availability

All data needed to evaluate the conclusions in the paper are present in the main text and/or the Supplementary Information. The master spreadsheet for the full proteomics and phosphoproteomics analysis is given in **Dataset S5**. Uniprot IDs mapped to EnsemblProtist IDs (v3 *P. tricornutum* genome) are given in **Dataset S6**. The mass spectrometry proteomics data have been deposited to the ProteomeXchange Consortium via the PRIDE partner repository with the dataset identifier PXD078910.

