## Supplementary Information for "Flexible phosphorus allocation to the phosphoproteome underpins diatom regulatory adaptations to fluctuating phosphate supply"

**This PDF includes**

Supplementary Figures S1-S12

Legend for Supplementary Video S1

Legends for Supplementary Datasets: S1-S6

Supplementary Materials and Methods

**Other supporting materials for this manuscript include the following:**

Video S1

Datasets S1-S6

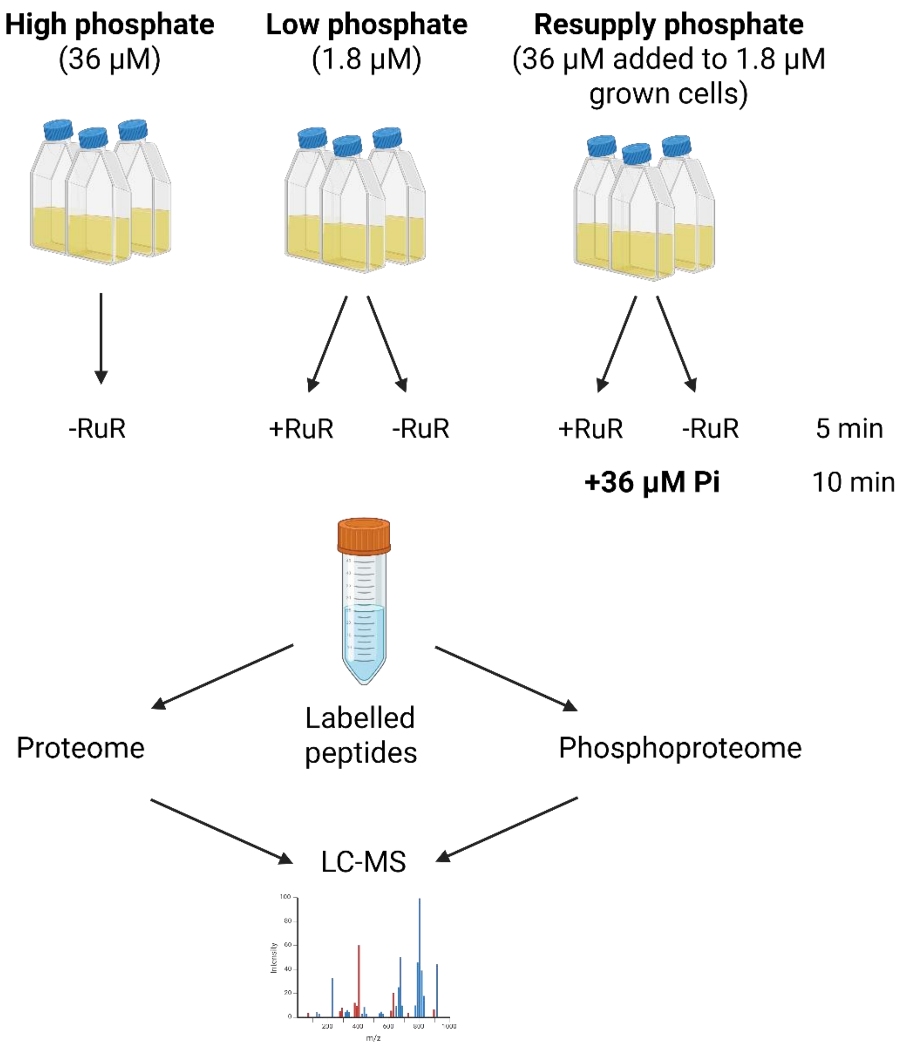


**Figure S1. Overview of the proteomics and phosphoproteomics experiment. A.** Schematic diagram summarising the experimental design for sample preparation. *Phaeodactylum tricornutum* cells were grown in high (36 µM), or low (1.8 µM) phosphate for four days. For the resupply treatments, 1.8 µM grown cells were resupplied with 36 µM phosphate and all cultures harvested 10 mins later**.** Cells treated with 5 µM ruthenium red (RuR) were incubated for 5 min prior to phosphate resupply for phosphate resupply cultures, or for 15 mins +RuR without phosphate resupply for the low phosphate +RuR treatment. Created with BioRender.com.

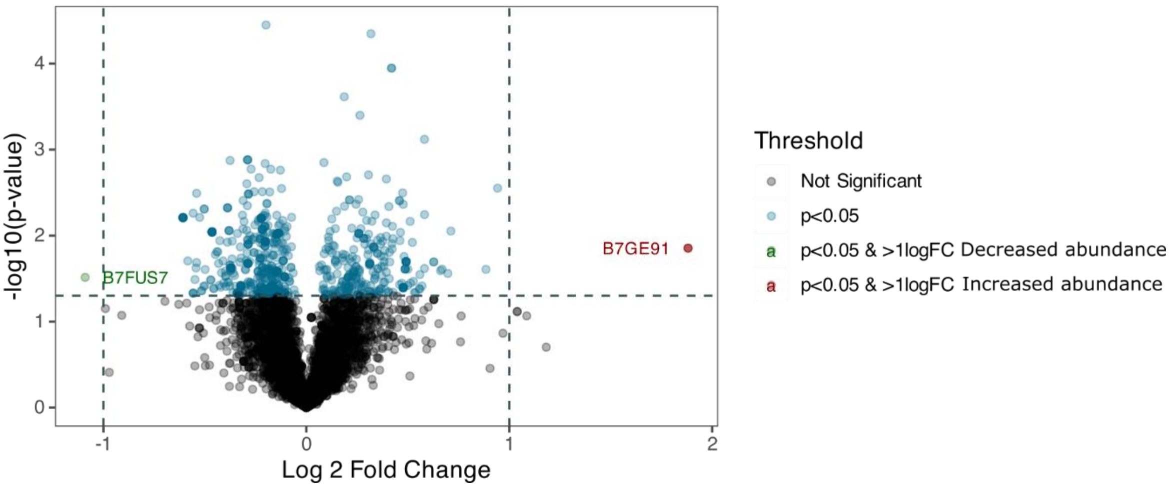


**Figure S2. Differential protein abundance of phosphate resupply versus phosphate treated *Phaeodactylum tricornutum*.** Cells were grown in low (1.8 mM phosphate for four days) and then resupplied with phosphate (36 mM) for 10 min before harvesting. The X-axis is the $log_2$fold change ($log_2$FC) of phosphate resupply versus low phosphate cells and the Y-axis is the negative log10 of the *p*-value for paired t-tests between the conditions described (*n*=3). Grey dots: not significant, blue dots: *p*-value<0.05, green dots: decrease in protein abundance with a $log_2$FC< -1 and a *p*-value <0.05, red dots: increase in protein abundance in phosphate resupply compared to low phosphate with a $log_2$FC>1 and a *p*-value<0.05. For phosphosites coloured green or red protein Uniprot accession numbers are given.

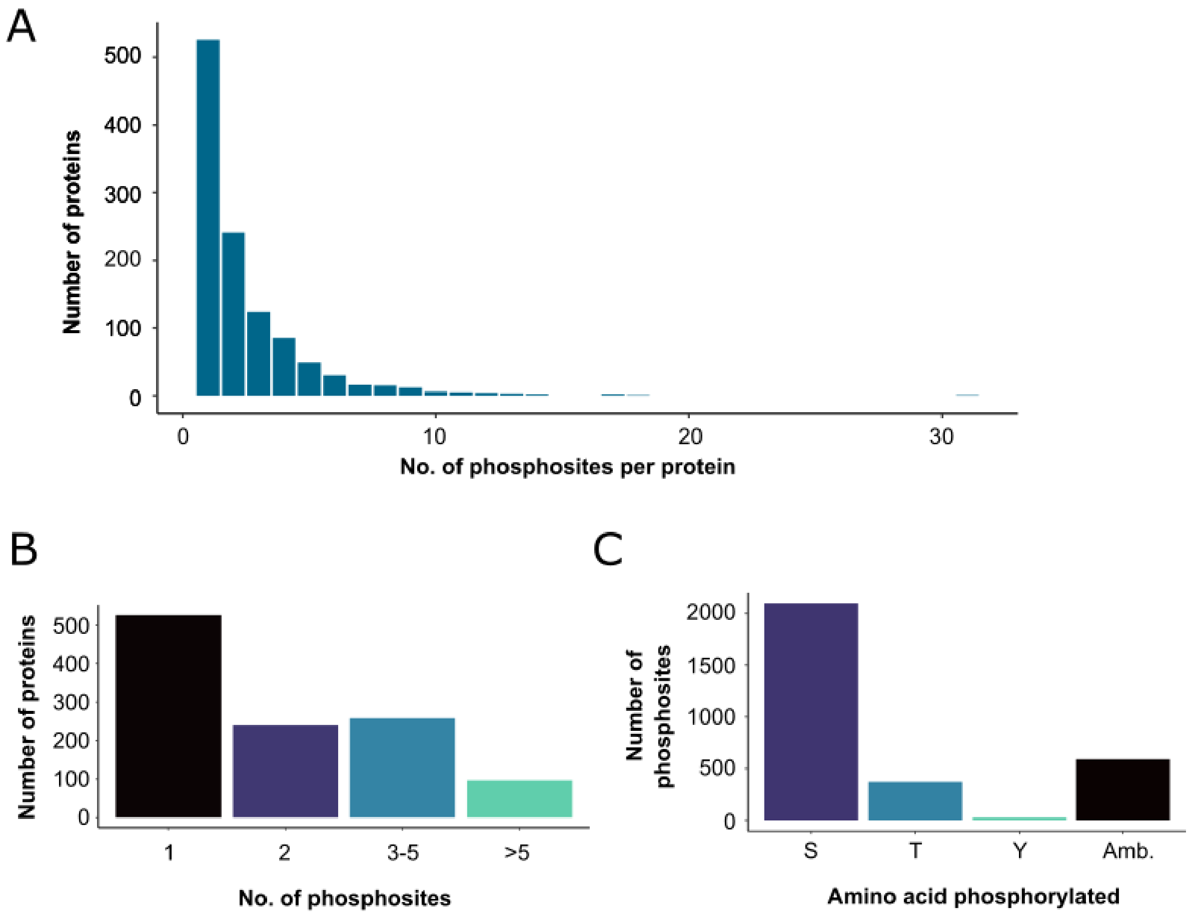


**Figure S3. Overview of *Phaoedactylum tricornutum* phosphoproteome from this study. A.** The distribution of phosphosites per protein. **B.** Number of phosphosites detected per protein. **C.** Number of phosphosites with phosphorylated serine (S), threonine (T), tyrosine (Y) and ambiguous (Amb.). The data is plotted for 3088 phosphosites detected on 1122 unique proteins that have corresponding protein abundance data available.

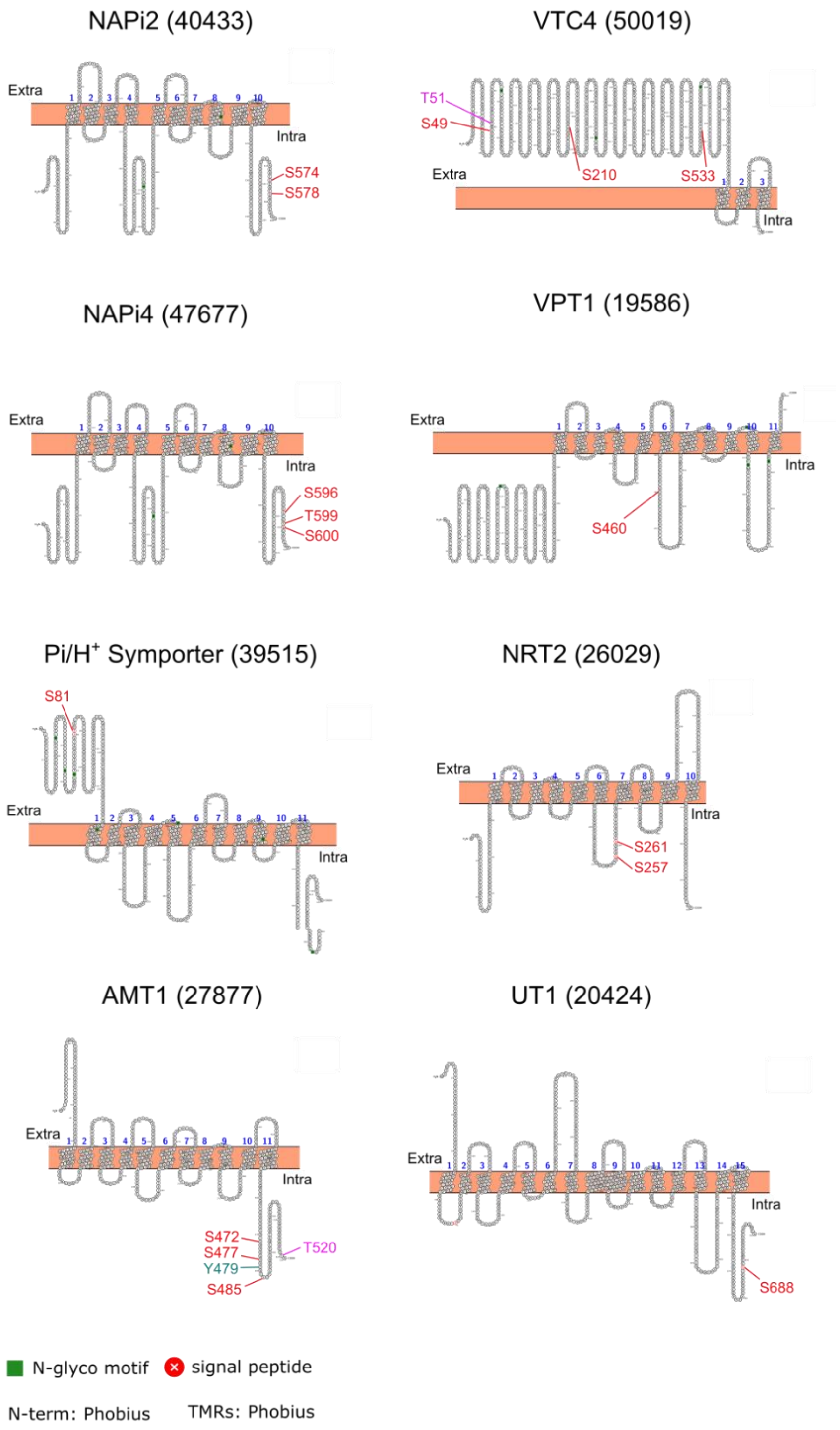


**Figure S4. Membrane topology and phosphosites of proteins containing transmembrane domains relating to P and N transport.** Protein topology as visualised using PROTTER (Omasits et al., 2014). Only phosphosites where the location is fully resolved are shown. Abbreviations for protein names are as follows: $Na^+$/phosphate transporters NAPi4 and NAPi2, vacuolar transport chaperone 4 (VTC4), vacuolar phosphate transporter (VPT1), nitrate transporter (NRT2), ammonium transporter 1 (AMT1), and urea transporter 1 (UT1).

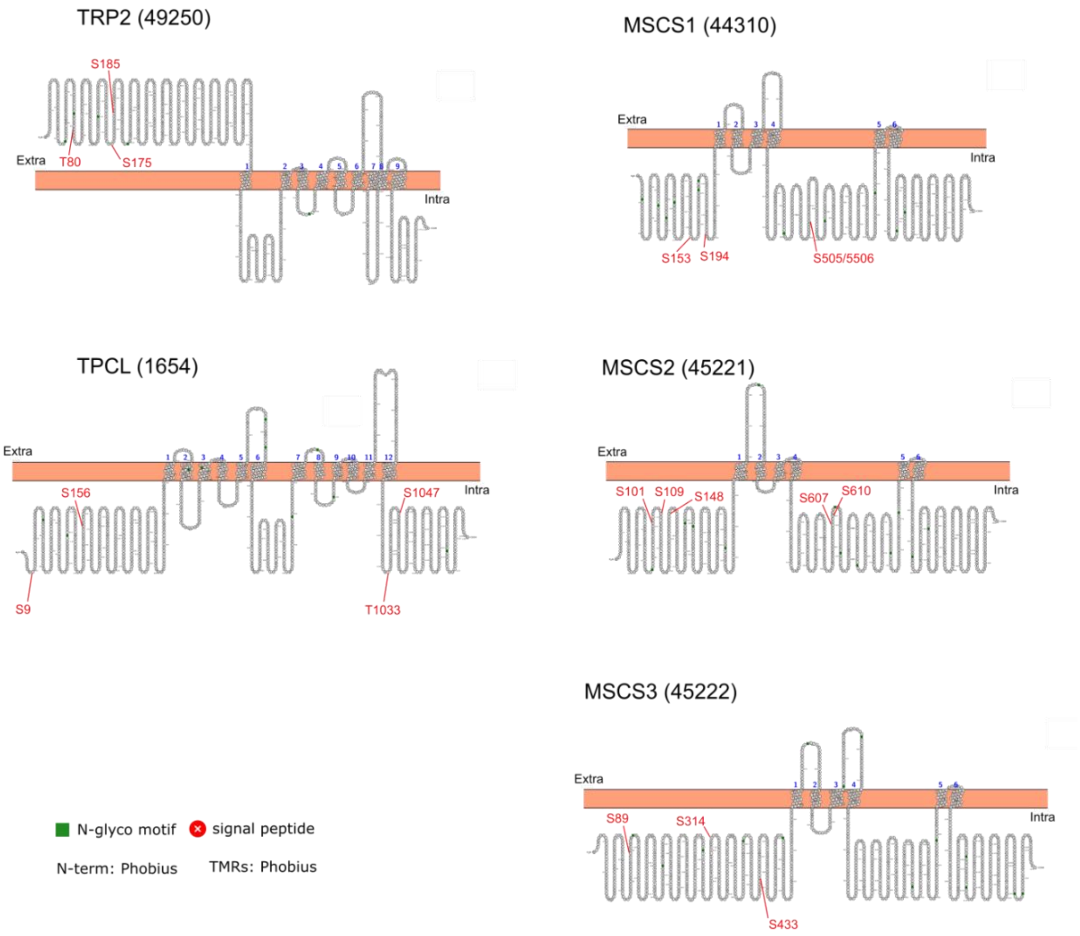


**Figure S5. Membrane topology and phosphosites of proteins containing transmembrane domains relating to $Ca^{2+}$ signalling.** Protein topology as visualised using PROTTER (Omasits et al., 2014), Only phosphosites where the location is fully resolved are shown. Abbreviations for protein names are as follows: transient receptor potential 2 (TRP2) channel, two-pore cation (TPC)-like channel, and mechanosensitive (MSCS1- MSCS3).

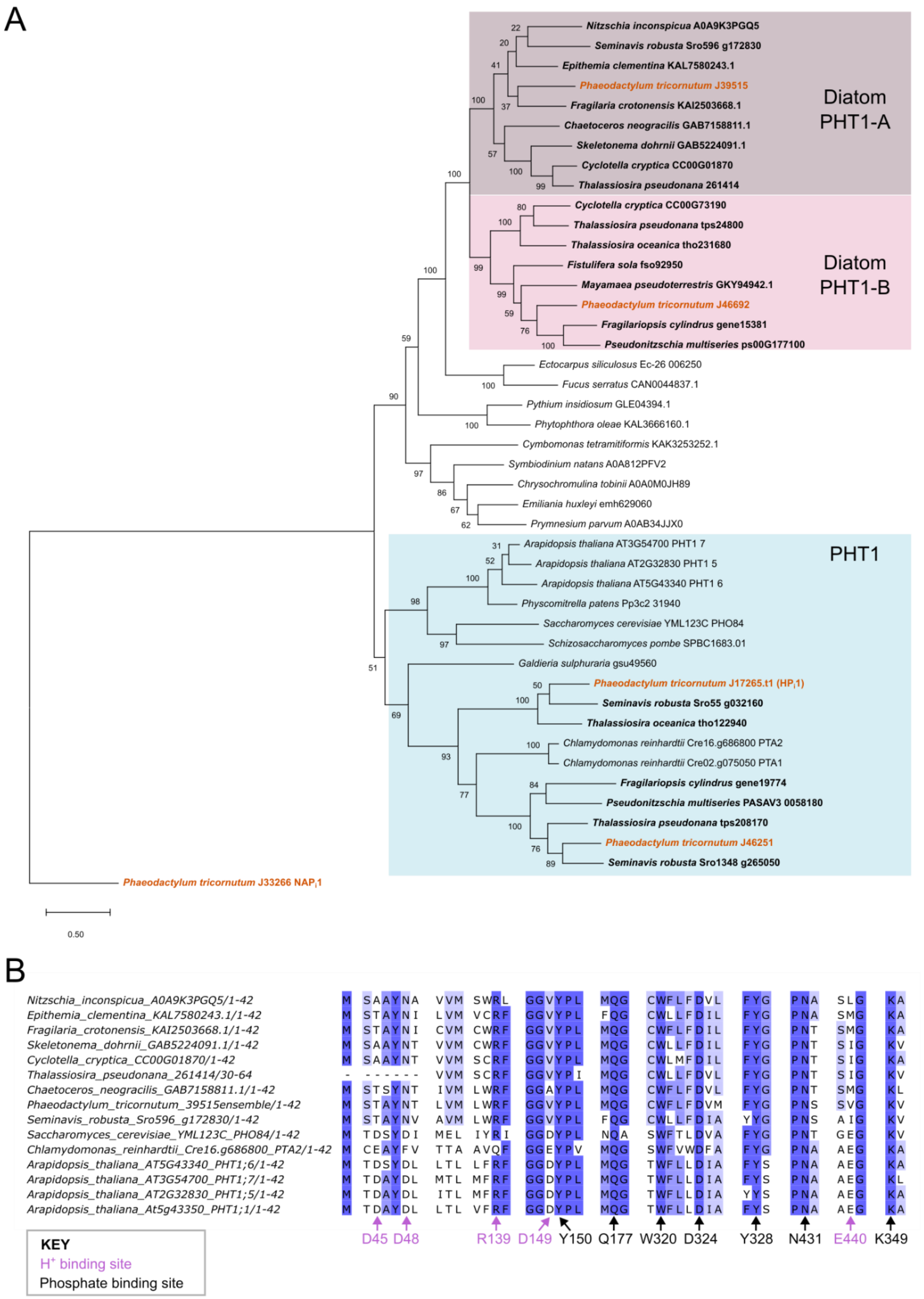


**Figure S6. *Phaeodactylum tricornutum* Pt39515 belongs to a novel diatom clade (PHT1-A) of intracellular phosphate transporters. A.** A maximum likelihood phylogenetic tree of Pt39515

compared to plant, algal and fungal $H^+$/phosphate transporters. The tree was reconstructed with 100 bootstraps using a LG (+Freq) model. The final alignment was 436 amino acids. **B**. A multiple sequence alignment showing percentage identity of diatom PHT1-A clade transporters compared to plant/green algal PHT1/PTA2 sequences and *Saccharomyces cerevisiae* PHO84, generated using Jalview 2.11.5.1. Conserved residues for $H^+$ and phosphate transport are labelled, based on the crystal structure and sequence of fungus *Piriformospora indica*, PiPT [54,55].

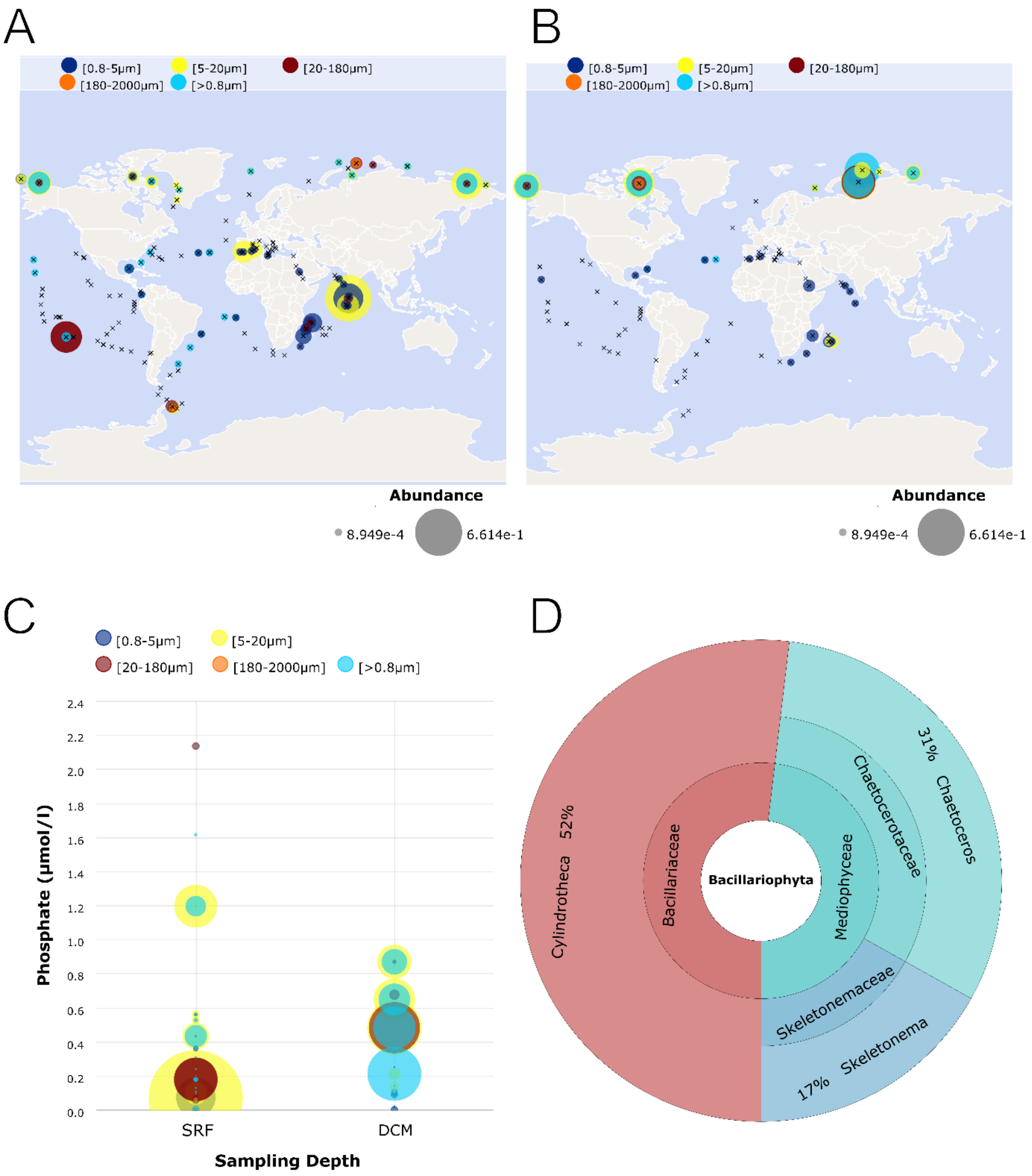


**Figure S7**. **Distribution of Pt39515 homologues in *TARA Oceans* Metagenome and Single Cell Assembled Genomes (SMAGs).** Geographic distribution of Pt39515 hits in **A.** the surface layer (SRF) and **B**. deep chlorophyll maximum (DCM). A strict cut-off e-value < $e^{-100}$ was used to query the database. **C.** Abundance of Pt39515 homologues with respect to phosphate (μml/l) levels. The taxonomic assignment of diatom hits is also given (**D**)

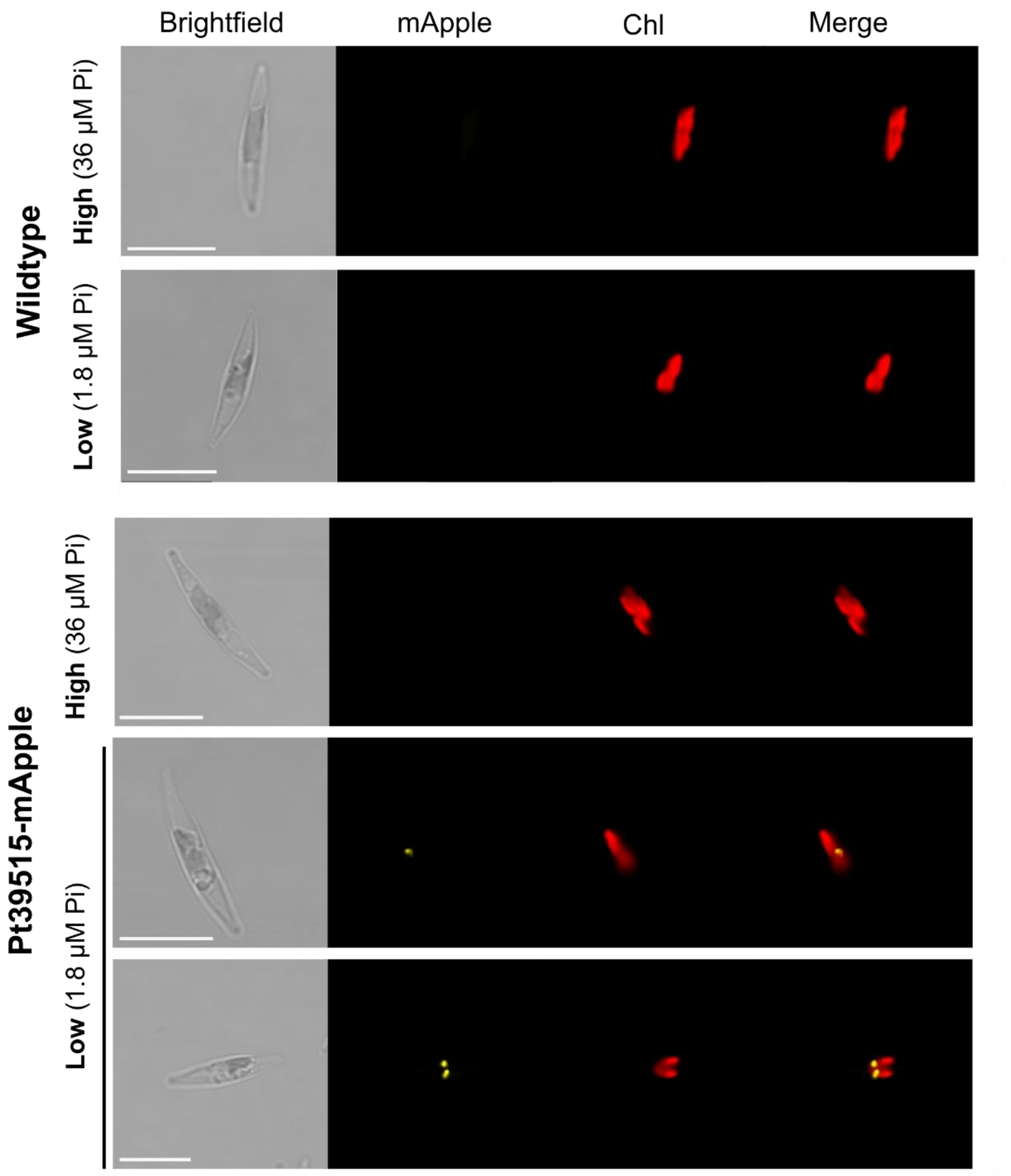


**Figure S8. Confocal microscope images of wildtype and Pt39515-mApple *P. tricornutum* cells.** Example images are given for cells grown with 36 µM (high) or 1.8 µM (low) phosphate conditions for four days. Scale bar: 10 µm.

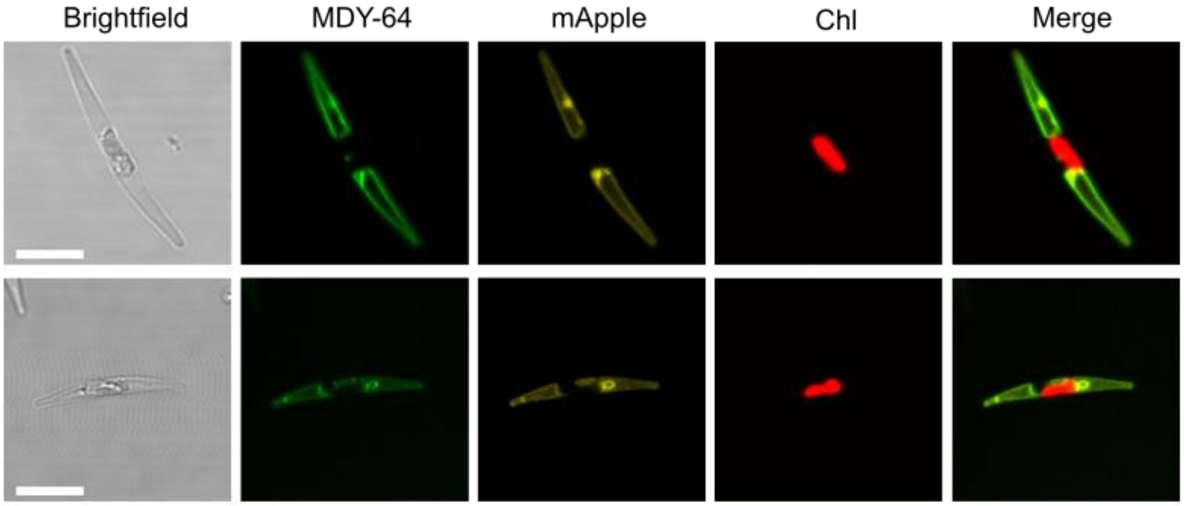


**Figure S9. Confocal microscope images of PtCDPK3-mApple cells co-stained with vacuole dye MDY-64**. Cells were grown 1.8 µM (low) phosphate conditions for four days. Two example images in cells grown in these conditions are given. Scale bar: 10 µm.

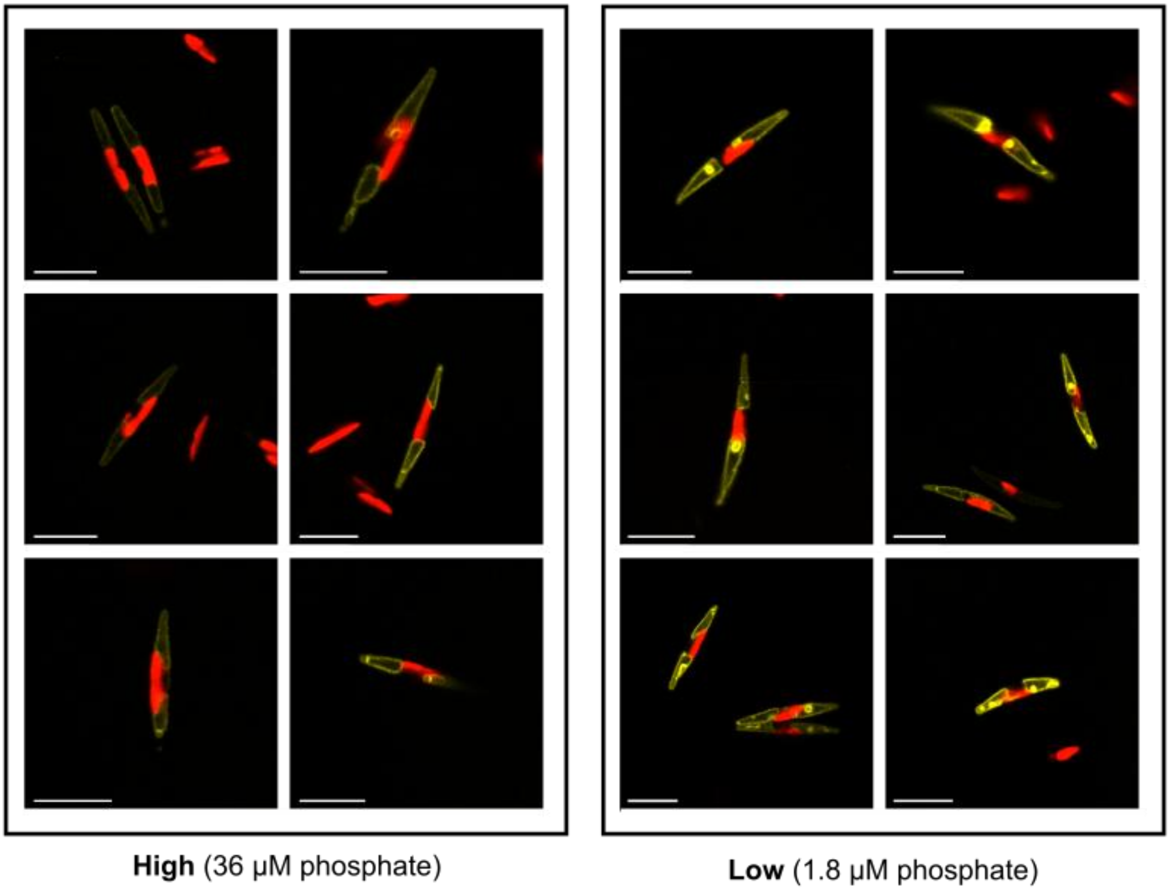


**Figure S10. Representative images of PtCDPK3-mApple cells grown in low and high phosphate conditions.** Cells were grown for four days in 36 μM (high) or 1.8 μM (low) phosphate conditions. Six example images for each condition are given. Scale bars: 10 μm.

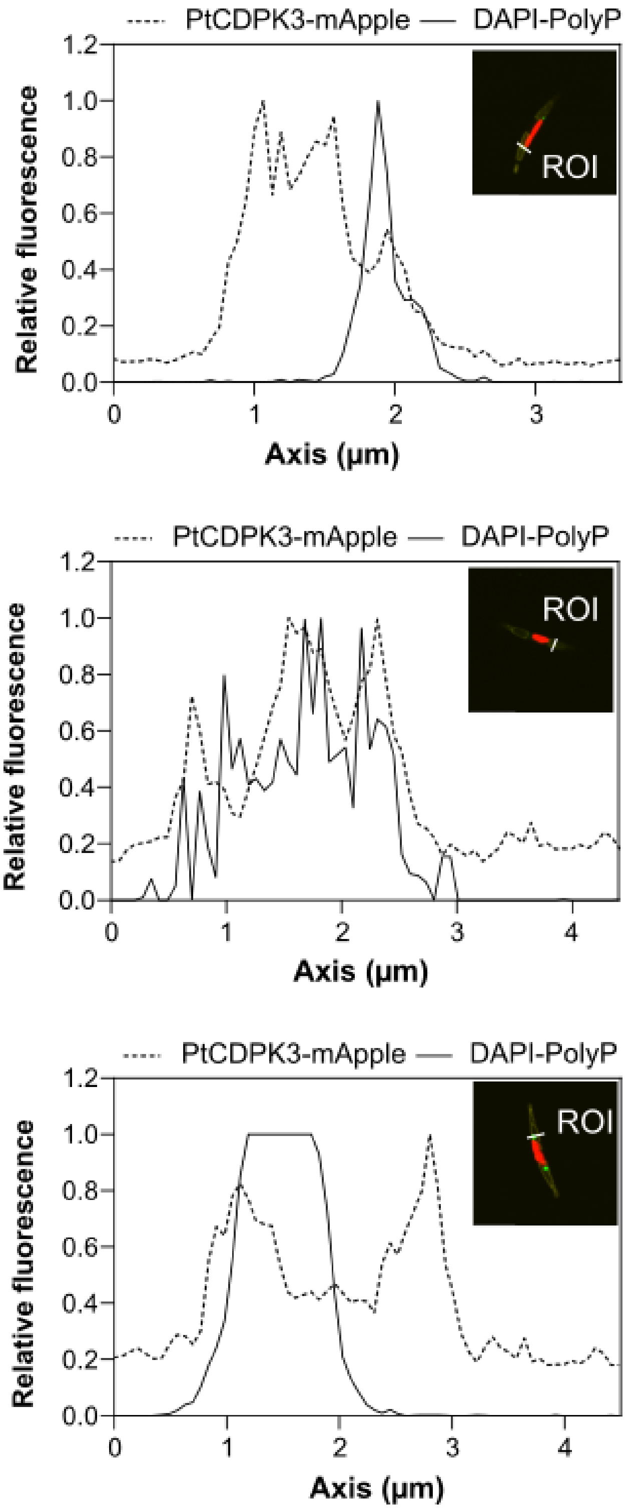


**Figure S11**. **PtCDPK3-mApple signal in relation to DAPI-stained polyP granules**. Representative relative fluorescence traces for PtCDPK3-mApple and DAPI-PolyP signal for the linear regions of interest (ROI) of images shown in Figure 6E including cells grown in 36 µM phosphate (top), 1.8 µM phosphate (middle) and 4 h phosphate resupply (bottom).

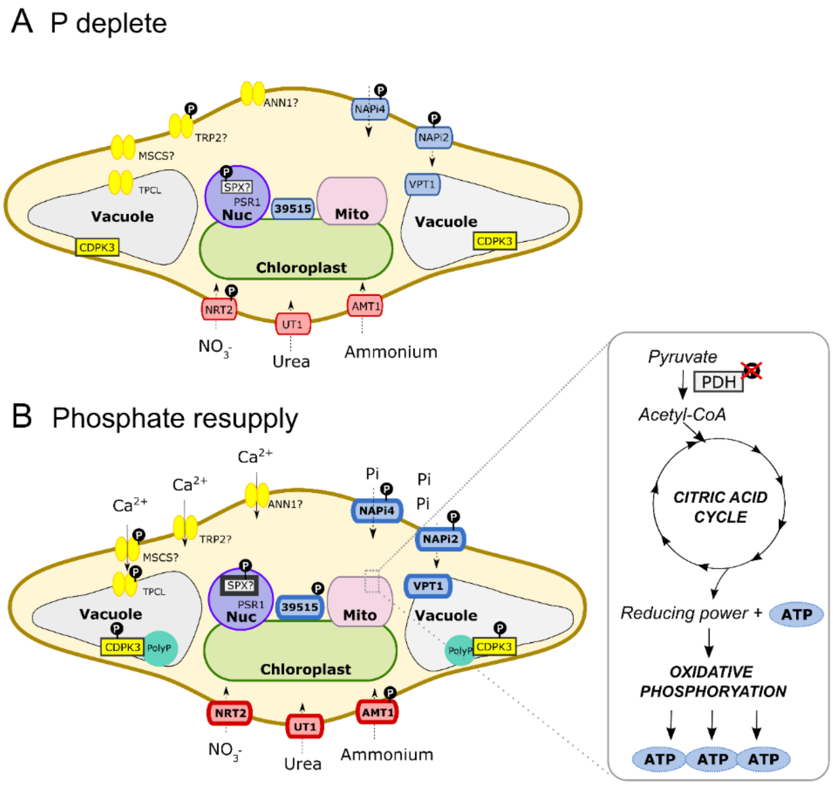


**Figure S12. Summary of changes in protein phosphorylation status of nitrogen, phosphorus and $Ca^{2+}$ signalling related machinery in response to (A) P limitation and (B) resupply.** Abbreviations and protein names are as follows: nitrate transporter (NRT2), urea transporter (UT1), ammonium transporter (AMT1), two-pore cation (TPC)-like channel, mechanosensitive (MSCS), $Ca^{2+}$-dependent protein kinase (CDPK), transient receptor Potential (TRP) channel, annexin (ANN1), vacuolar phosphate transporter (VPT1), pyruvate dehydrogenase (PDH), phosphorus starvation response 1 (PSR1), phosphate-$H^+$ symporter (39515), SYG1, Pho81 and XPR1 (SPX) domain protein, and $Na^+$/phosphate transporters NAPi4 and NAPi2.

**Legend for Supplementary Video S1**

**Video S1.** Video of 6-day old low phosphate (1.8 µM) grown PtCDPK3-mVenus cell, over a time course of 3 hrs following 36 µM phosphate resupply, imaging cells every 10 minutes.

**Legends for Supplementary Datasets S1-S6**

**Dataset S1.** A list of proteins showing significantly (p-value<0.05) altered abundance in low versus high phosphate conditions (including those showing a $log_2$fold change of less than -1 or greater than 1). Protein identifiers, annotations as well as results of a t-test performed on the total - $Log_2$ normalised abundances of proteins in the conditions are listed.

**Dataset S2.** A list of phosphosites showing significantly (p-value<0.05) altered phosphorylation in low versus high phosphate conditions. Protein identifiers, annotations and phosphosite metrics as well as Phos - Adjusted $Log_2$ Normalised Abundance values are provided. Only those phosphosites showing a $log_2$fold change of less than -1 or greater than 1 are given.

**Dataset S3.** Analysis of the impact of ruthenium red (RuR) on phosphosites showing significantly altered phosphorylation status in resupply versus low P conditions. All phosphosites showing significantly altered phosphorylation status (based on Phos - Adjusted $Log_2$ Normalised Abundance) in the P resupply versus low P conditions are shown (purple). Corresponding data for resupply+RuR versus resupply-RuR (blue), and low+RuR versus low-RuR conditions (grey) are also given, indicating significance thresholds. Protein identifiers, annotations and phosphosite metrics are provided.

**Dataset S4.** Protein identifiers, sequence information and predicted localisation of $H^+$/phosphate transporters shown in Figure S6B. Protein localisation prediction results from DeepLoc2.1, HECTAR, ASAFIND2 and TargetP2.0 prediction software are given.

**Dataset S5.** A list of all proteins and phosphosites across all treatments with selected pairwise comparisons provided. A guide for filtering and interpreting the data is given in the Dataset S5_Guide tab. Phosphosite location predictions were based on the *P. tricornutum* (strain CCAP1055/1) Uniprot database (downloaded June 2022: 10465 entries). Uniprot IDs mapped to EnsemblProtist IDs (v3 *P. tricornutum* genome) are given in Dataset S6.

**Dataset S6.** Uniprot IDs mapped to EnsemblProtist IDs (v3 *Phaeodactylum tricornutum* genome).

### Supplementary Materials and Methods

#### Sample preparation for quantitative proteomics

Cells were inoculated to a starting cell density of 3 x $10^4$ cell/ml in 100 ml flasks, and cultures grown in triplicate for four days in either high (36 µM), low (1.8 µM) or phosphate resupply treatments (i.e. cells grown as in the low phosphate condition ready for resupply after four days). On day four, cells grown in low P conditions (both for low and resupply treatments) were split in two. One batch was incubated without (-RuR) and another with (+RuR) ruthenium red to a final concentration of 5 µM for 5 mins (**Figure S1**). Phosphate (36 µM) was resupplied to the resupply treatments, and 10 mins later, all treatments were harvested by centrifugation at 4°C for 10 mins 4000 x g and snap frozen in liquid $N_2$ and stored at -80°C until further processing.

For total protein analyses, cell pellets were re-suspended in 50 µl or 100 µl of RIPA lysis and extraction buffer (Thermo-Fisher Scientific, UK) for phosphate deplete and replete cells, respectively. 100x Halt protease inhibitor (cOmplete™ protease inhibitor cocktail, Merck, Germany) was added to a final concentration of 1x. Additionally, 1 tablet of phosSTOP™ phosphatase inhibitor (Roche, Switzerland) per 50 ml buffer was used. Samples were vortexed for 30 s and sonicated for 5 mins in a sonication bath with ice. The samples were then centrifuged for 15 mins at 11,000 x g at 4°C and supernatants were transferred to clean tubes. Total protein was then quantified using a Pierce BCA Protein Assay kit (Thermo-Fisher Scientific, UK), according to manufacturer's instructions and assessed by sodium dodecyl sulfate–polyacrylamide gel electrophoresis (SDS-PAGE) according to manufacturer's instructions (Sigma Aldrich, UK).

#### TMT Labelling, high pH reversed-phase chromatography and phospho-peptide enrichment.

Aliquots of 100 µg of each sample were digested with trypsin (2.5 µg trypsin per 100 µg protein; 37°C, overnight), labelled with Tandem Mass Tag (TMTpro) sixteen plex reagents according to the manufacturer's protocol (Thermo-Fisher Scientific, UK) and the labelled samples pooled.

For the total proteome analysis, an aliquot of 50 µg of the pooled sample was desalted using a SepPak cartridge according to the manufacturer's instructions (Waters, Milford, Massachusetts, USA). Eluate from the SepPak cartridge was evaporated to dryness and resuspended in buffer A (20 mM ammonium hydroxide, pH 10) prior to fractionation by high pH reversed-phase chromatography using an Ultimate 3000 liquid chromatography system (Thermo-Fisher Scientific, UK). In brief, samples were loaded onto an XBridge BEH C18 Column (130Å, 3.5 µm, 2.1 mm X 150 mm, Waters, UK) in buffer A and peptides eluted with an increasing gradient of buffer B (20 mM Ammonium Hydroxide in acetonitrile, pH 10) from 0-95% over 60 minutes. The resulting fractions (concatenated into 20 in total) were evaporated to dryness and resuspended in 1% formic acid prior to analysis by nano-LC MS/MS using an Orbitrap Fusion Lumos mass spectrometer (Thermo-Fisher Scientific, UK).

For the phosphoproteome analysis, the remainder of the TMT-labelled pooled sample was also desalted using a SepPak cartridge (Waters, Milford, Massachusetts, USA). Eluate from the SepPak cartridge was evaporated to dryness and subjected to $TiO_2$-based phosphopeptide enrichment according to the manufacturer's instructions (Pierce Biotechnology, Rockford, USA). The flow-through and washes from the $TiO_2$-based enrichment were then subjected to FeNTA-based phosphopeptide enrichment according to the manufacturer's instructions (Pierce Biotechnology). The phospho-enriched samples were again evaporated to dryness and then resuspended in 1% formic acid prior to analysis by nano-LC MSMS using an Orbitrap Fusion Lumos mass spectrometer (Thermo-Fisher Scientific, UK).

**Nano-LC Mass Spectrometry**

High pH reversed phase fractions (total proteome analysis) or the phospho-enriched fractions (phospho-proteome analysis) were further fractionated using an Ultimate 3000 nano-LC system in line with an Orbitrap Fusion Lumos mass spectrometer (Thermo-Fisher Scientific, UK). In brief, peptides in 1% (vol/vol) formic acid were injected onto an Acclaim PepMap C18 nano-trap column (Thermo-Fisher Scientific, UK). After washing with 0.5% (vol/vol) acetonitrile 0.1% (vol/vol) formic acid peptides were resolved on a 250 mm × 75 μM Acclaim PepMap C18 reverse phase analytical column (Thermo-Fisher Scientific, UK) over a 150 mins organic gradient, using 7 gradient segments (1-6% solvent B over 1 min, 6-15% B over 58 min, 15-32% B over 58 min, 32-40% B over 5 min, 40-90% B over 1 min, held at 90% B for 6 mins and then reduced to 1% B over 1 min) with a flow rate of 300 nL $min^{-1}$. Solvent A was 0.1% formic acid, and Solvent B was aqueous 80% acetonitrile in 0.1% formic acid. Peptides were ionized by nano-electrospray ionization at 2.0kV using a stainless-steel emitter with an internal diameter of 30 μm (Thermo-Fisher Scientific, UK) and a capillary temperature of 300°C.

All spectra were acquired using an Orbitrap Fusion Lumos mass spectrometer controlled by Xcalibur 3.0 software (Thermo-Fisher Scientific, UK) and operated in data-dependent acquisition mode using an SPS-MS3 workflow. FTMS1 spectra were collected at a resolution of 120000, with an automatic gain control (AGC) target of 200000 and a max injection time of 50 ms. Precursors were filtered with an intensity threshold of 5000, according to charge state (to include charge states 2-7) and with monoisotopic peak determination set to Peptide. Previously interrogated precursors were excluded using a dynamic window (60s +/-10 ppm). The MS2 precursors were isolated with a quadrupole isolation window of 0.7 m/z. ITMS2 spectra were collected with an AGC target of 10000, max injection time of 70 ms and CID collision energy of 35%.

For FTMS3 analysis, the Orbitrap was operated at 50000 resolutions with an AGC target of 50 000 and a max injection time of 105 ms. Precursors were fragmented by high energy collision dissociation (HCD) at a normalised collision energy of 60% to ensure maximal TMT reporter ion yield. Synchronous Precursor Selection (SPS) was enabled to include up to 10 MS2 fragment ions in the FTMS3 scan.

**Proteomics data analysis**

The raw data files were processed and quantified using Proteome Discoverer software v2.4 (Thermo-Fisher Scientific, UK) and searched against the UniProt *P. tricornutum* (strain CCAP1055/1) database (downloaded June 2022: 10465 entries) using the SEQUEST HT algorithm. Peptide precursor mass tolerance was set at 10 ppm, and MS/MS tolerance was set at 0.6 Da. Search criteria included oxidation of methionine (+15.995 Da), acetylation of the protein N-terminus (+42.011 Da) and methionine loss plus acetylation of the protein N-terminus (-89.03 Da) as variable modifications and carbamidomethylation of cysteine (+57.0214) and the addition of the TMTpro mass tag (+304.207) to peptide N-termini and lysine as fixed modifications. For the phospho-proteome analysis, phosphorylation of serine, threonine and tyrosine (+79.966) was also included as a variable modification. Searches were performed with full tryptic digestion and a maximum of 2 missed cleavages were allowed. The reverse database search option was enabled, and all data was filtered to satisfy false discovery rate (FDR) of <5%, with 90.7% of proteins identified with a <1% FDR.

**Protein and phosphopeptide abundance processing**

Protein abundances were normalised within each sample to total peptide amount before being $Log_2$ transformed. Phosphorylation status of identified peptide spectral matches (PSMs) was determined by Proteome Discover Software (PD2.1) and the site of phosphorylation predicted using the PhosphoRS module. Phosphorylation sites predicted with greater than 70% confidence were taken as the likely phosphorylation site, and phosphopeptides identified with identical sequences and predicted phosphorylation sites, were combined to provide improved quantitation and confidence. Phosphosite location predictions were based on the *P. tricornutum* (strain CCAP1055/1) Uniprot database (downloaded June 2022: 10465 entries). If a peptide was predicted to be phosphorylated (based on its mass), but the software was unable to assign the site, the site was listed as “Ambiguous”. Where multiple phosphorylation events are unable to be located to specific sites within a peptide, the word “Ambiguous” is repeated the corresponding number of times. As peptides can often be matched to multiple proteins, the list of proteins to which each peptide matched was searched against the list of master proteins in the total protein analysis, and if a matching protein was identified, this protein was used as the master protein for that peptide. Normalisation of phosphopeptide abundance was performed using the normalisation factor generated from analysis of the corresponding samples in the total protein dataset. For this, $log_2$ normalised protein abundance was subtracted from the $log_2$ normalised phosphopeptide abundance to adjust the phosphopeptide abundance for any changes in the total protein and yield a $log_2$ normalised phosphopeptide abundance value.

Statistical significance was determined using paired t-tests between the conditions of interest to determine p-values. Data was analysed using R Studio statistical software (v4.2.2).

**Experiments quantifying cellular ATP levels of *P. tricornutum* under different P regimes.**
Wildtype *P. tricornutum* cultures were inoculated to a starting density of 3 x $10^4$ cells $ml^{-1}$ and grown in triplicate in 300 ml of modified f/2 medium -Si and vitamins. Phosphate was added according to the different experimental P treatments, as follows: high (36 µM initial phosphate), low (1.8 µM initial phosphate) or phosphate resupply (1.8 µM phosphate grown cells resupplied with 36 µM phosphate on day 6) treatments. Fifty ml of each sample was harvested prior to phosphate resupply (to the resupply treatment) i.e. t=0. Subsequent samples were taken 1, 2, and 4 h post resupply for the low and resupply treatments. Cell counts were performed at t=0 using a Beckman Coulter Counter (Beckman Coulter, UK) with a 3-9 µm aperture. Samples were resuspended in 200 or 1200 µl (for low/resupply and high grown cells, respectively) of ReadiUse™ Yeast Cell Lysis Buffer (catalogue no. 24102, AAT Bioquest, USA) with glass beads (0.1–1.5 mm, autoclaved). Cells were lysed by vortexing (5 x 15 secs, alternated with 2 mins chilling on ice). For ATP quantification a PhosphoWorks™ Fluorimetric ATP Assay Kit with minimal interference from ADP/AMP (catalog no. 21620, AAT Bioquest, USA), was employed following the manufacturer's protocol using an ATP standard curve for calibration. Standards and experimental samples (50 µl) were pipetted into a black 96-well plate (catalog no. 655209 Greiner) and 50 µl ATP working solution added. The samples were left to incubate at room temperature for 30 mins in the dark before fluorescence was measured on a spectrophotometer plate reader at Ex/Em=540/590 nm (Cutoff = 570 nm). Quantified ATP levels were normalised per cell for each culture.

**Localisation of Pt39515 and PtCDPK3 in *P. tricornutum***

The *P. tricornutum Pt39515* and *PtCDPK3* genes, including their upstream promoter region (approx. 700 bp 5' of the translation initiation site) but excluding the stop codon, were amplified from 10 ng *P. tricornutum* genomic DNA (extracted using the DNA isolation procedure described by Newman et al., (1990) using Q5 High Fidelity DNA Polymerase (New England Biolabs). Primer sequences for *Pt39515* were as follows: Keh405F atagaaccagatcccccgggctgcaAAGTACCTCCTGCTACGGGT and Keh406R gttttcttcacccttggaaaccataggATCATCGTAGTAGTCAAAGAGCGAG, and the following thermocycling conditions used: 95°C (pre-heated block), 98°C 30 sec, 33 x (98°C 5 sec, 67°C 10 sec, 72°C 90 sec), 72°C for 2 min. For *PtCDPK3*, primers Keh403F atagaaccagatcccccgggctgcaGTGTCTGTCGGCCGGAATTA and Keh404R gttttcttcacccttggaaaccataggCGCAGGAATGTCTTCTACTTCC, were used with the following PCR conditions: 95°C (pre-heated block), 98°C 30 sec, 33 x (98°C 5 sec, 66°C 10 sec, 72°C 90 sec), 72°C for 2 min. The amplicons were cloned into pPhat-1_mApple vector (a derivative of the pPHA-T1 (accession AF219942) adapted as described by Helliwell et al., (2019) containing a codon optimised mApple gene (protein id. WYD57316.1) and BlastR). Genes were cloned into the vector via *StuI* and *PstI* sites, using the NEBuilder HiFi DNA Assembly Cloning Kit (New England Biolabs), to create mApple fusions. The resulting constructs were used to transform *P. tricornutum* by biolistic bombardment as described previously (62) with blasticidin (8 ug/ml) selection (62). Transformants were screened for fluorescence with a Leica SP8 confocal microscope using a 63x oil objective, with mApple fluorescence excited at 561 nm and emission measured at 600 ± 20 nm.

**MDY-64 staining of *P. tricornutum* vacuolar membranes**

*P. tricornutum* vacuolar membranes were stained using the dye MDY-64 (Thermo Fisher Scientific, UK). 2.5 µL of 0.1 mM MDY-64 diluted in 100% DMSO was added to 500 µl of WT and PtCDPK3-mApple *P. tricornutum* cells in f/2 -Si liquid culture on day four post sub-culturing. Cells were gently mixed through pipetting and transferred immediately to 0.1% poly-L-lysine coated glass-bottomed dishes for imaging on a Leica TCS SP8 confocal microscope with 458 nm excitation and a 510 ± 25 nm emission. mApple fluorescence was excited at 561 nm and emission measured at 600 ± 20 nm. Chlorophyll was excited at 488 nm and the emission wavelength 700 ± 50 nm.

**DAPI staining of *P. tricornutum* PolyP**

DAPI (Thermo Fisher Scientific, UK) was used to stain *P. tricornutum* DNA and polyphosphate (PolyP). 4 µl of 1 mM DAPI diluted in PBS was added to 200 µl (final concentration of 20 µM) of *P. tricornutum* cells in f/2 -Si liquid culture on day four or five post sub-culturing. Cells were gently mixed through pipetting and left to incubate in the dark for two hours. Cells were subsequently washed via centrifugation for one min at 13,000 x g, liquid was removed via pipetting and cells were resuspended in fresh f/2 – Si liquid media. This wash procedure was repeated three times and cells were then transferred to 0.1% poly-L-lysine coated glass-bottomed petri dishes for imaging on a Leica TCS SP8 confocal microscope. A two-sequence imaging protocol was used to first image DAPI through laser excitation at a wavelength of 405 nm and an emission detection of 535 – 570 nm for PolyP. The second sequence excited mApple and chlorophyll fluorescence, as described above.
